# Metagenomic co-assembly reveals mobile antibiotic resistance genes in airborne microbiomes in the Eastern Mediterranean

**DOI:** 10.1101/2024.09.26.615105

**Authors:** Burak Adnan Erkorkmaz, David Zeevi, Yinon Rudich

## Abstract

Antibiotic resistance is a significant threat to global ecological health. Mobile resistance genes are of particular concern due to their ability to transfer between organisms. In environmental settings and within the microbiota of humans and animals, these genes can sometime move into pathogens, promoting the spread of resistance. While various dispersal mechanisms have been studied, the role of air masses and dust storms in spreading antibiotic resistance genes globally remains largely unexplored. We analyzed air samples from non-dusty conditions and Middle Eastern dust storms using a metagenomic co-assembly approach to identify mobile antibiotic resistance genes dispersed by these air masses. Our analysis shows that co-assembling low-biomass air samples of similar origin produces longer genome fragments, allowing for a more detailed bioinformatic analysis that links functional features to specific taxa. Through this method, we identified genes associated with resistance to antibiotics such as aminoglycosides, beta-lactams, bleomycin, fosfomycin, chloramphenicol, daunorubicin, fluoroquinolones, glycopeptides, quinolones, streptomycin, and tetracyclines. Approximately 10% of these genes were linked to mobile genetic elements. These findings demonstrate the potential for global spread of resistance genes across ecosystems via air masses, particularly dust storms, and underscore the need for continuous atmospheric monitoring and further research to develop strategies that mitigate their impact on public health, the environment, and agriculture.

## Introduction

The atmosphere harbors a microbiome that plays an important role in the functioning of Earth’s ecosystems. These microorganisms can act as condensation and ice nuclei ^1-3^ and contribute to atmospheric chemical processes ^4,5^. However, compared to other habitats, the atmospheric microbiome remains relatively underexplored, particularly regarding its microbial characteristics and their ecological impacts. Environmental changes, such as temperature fluctuations and shifts in particulate matter (PM_10_) concentrations, can significantly influence the atmospheric microbiome ^6-8^. On days with lower PM_10_ concentrations, the atmospheric microbiome typically reflects the ecological traits of the local environment, with minimal contributions from distant sources. Conversely, during dusty conditions with high PM_10_ levels, the microbiome undergoes significant changes, primarily reflecting the ecological characteristics of the particulate matter’s origin ^7,9-11^. Previous studies have indicated that dust storms can disperse harmful biological agents such as potential pathogens and antibiotic resistance and virulence genes ^7,9,10,12-14^. These traits can originate from natural habitats or sources such as hospitals, wastewater treatment facilities, and agricultural or livestock operations treated with antibiotics ^15-19^. If these contaminants are released into the atmosphere in significant quantities, they may pose additional ecological and health risks ^20^.

The emerging field of bioaerosol studies has advanced significantly with the development of metagenomic tools. This technique allows for the direct analysis of nucleic acids, eliminating the biases associated with gene amplification commonly used in previous atmospheric microbiome studies. However, the low-biomass nature of the atmosphere often presents technical challenges ^21-23^. Accurate identification of functional characteristics of microbiomes requires tens of millions of high-quality sequencing reads, which can be difficult to obtain from atmospheric samples. These challenges complicate our understanding of the dynamics and the content of the atmospheric microbiome, especially during dust storms, where the sampling window to capture the storm’s peak is typically limited to a few hours. While extending the sampling duration can increase the collected biomass for a more comprehensive metagenomic analysis, this approach compromises the specificity of dust storm representation. Previous studies have shown that optimizing sampling and DNA extraction can improve taxonomic analysis, but it remains unclear how efficient this optimization is in connecting different aspects of the microbiome ^21,22,24,25^. Higher DNA recovery during extraction does not always guarantee deeper resolution in subsequent bioinformatic analyses. To connect the various functional characteristics of the microbiome, it is necessary to assemble genomic fragments that are sufficiently long to be informative for bioinformatic analysis.

Metagenomic assembly can be performed by individually assembling the reads from the same sample or by co-assembling reads from a group of samples. The choice of method should be tailored to specific research questions. The co-assembly approach offers the advantage of recovering low-abundance genes that may not be assembled from individual samples. Additionally, it enables the creation of longer contigs, facilitating more detailed analysis and the ability to connect microbial traits of interest ^26,27^. However, coassembly works best when samples are of similar origin, as this method may face challenges in diverse and low-biomass communities. In such cases, the total number of reads is lower, and these reads may come from ecologically distant taxa, making it difficult to generate accurate assemblies. Consequently, assembling algorithms might struggle to produce reliable results in these scenarios.

In our recent study, we demonstrated significant associations between dust storms, potential pathogens, and antibiotic and virulence-related genes using metagenomics. Our findings revealed that soil bacteria are the primary carriers of antibiotic and virulence-related traits. Only few DNA fragments (contigs) from pathogenic species contained these genes ^7^. This is likely due to the atmospheric microbiome’s remarkable diversity and low biomass, which result in highly fragmented genomes and make accurate analysis challenging. Consequently, we could not determine whether these genes were already mobile, as accurately identifying certain microbial aspects requires longer DNA reads. Mobile genetic elements are particularly important because they can indicate the potential mobility of genes and their associated functions, such as antibiotic resistance, within a community ^28,29^. This process is known as horizontal gene transfer. Therefore, profiling these elements and understanding their association with antibiotic resistance genes provide further insights into the environmental and public health relevance of these traits ^30^.

Hence, the primary objectives of this study are to evaluate the effectiveness of co-assembling atmospheric genetic data in addressing some of the previously discussed technical limitations and to determine whether antibiotic resistance traits are mobile within the atmospheric microbiome, particularly during dust storms, by analyzing longer DNA fragments. Additionally, we aim to investigate whether antibiotic resistance and virulence-related traits are associated with pathogenic species.

## Materials and Methods

### Sample Collection and Atmospheric Conditions during the Sampling Campaign

We conducted air sampling from the rooftop of a four-story building at the Weizmann Institute of Science in Rehovot, Israel (31.9070°N, 34.8102°E; 80 m above mean sea level) over a 64-day period from the end of March to the end of May 2022. We obtained particulate matter concentration (PM, μg m^−3^) and temperature (T, °C) data from the Rehovot Air Monitoring station, located approximately 1 km from the sampling site. Details on the origin of air masses, dust, and sample collection, as well as atmospheric conditions during the campaign, are provided in a recent study ^7^ and in Supporting Figure 1. During this period, we collected a total of 45 air samples.

**Figure 1.**
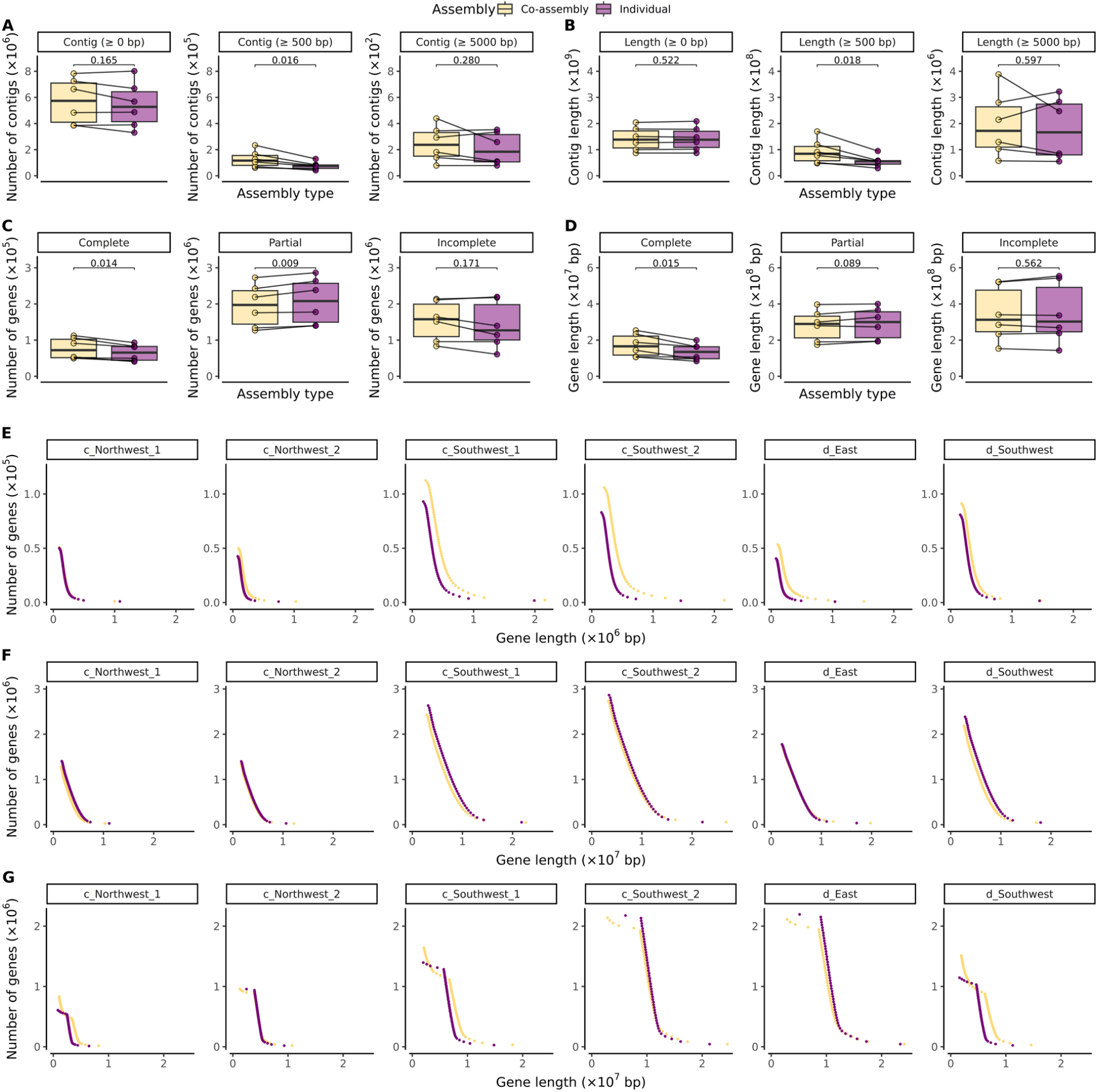
Comparison of co-assembly method performance to individual assembly. Differences in (A) total contig number, (B) total contig length, (C) total gene number and (D) total gene length between the two assembly approaches were assessed using a two-sided paired *t*-test. Boxplots display median values as center lines, with lower and upper hinges representing the 25^th^ and 75^th^ percentiles. Whiskers extend from the hinges to the smallest and largest values within 1.5 times the interquartile range (IQR), and outliers are shown as red dots. Cumulative distribution of gene lengths for the two assembly approaches for (E) complete, (F) partial and (G) incomplete genes in the two assembly approaches. For visual clarity, genes in each sample were divided into 50 bins, with each bin containing an equal number of genes sorted in descending order of length. Within each bin, total gene length and the number of genes were calculated.

For the metagenomic analysis, we utilized the Coriolis µ microbial air sampler (Bertin Technologies). Samples were collected for 2 hours at a flow rate of 200 L/min. This impinger captures particles larger than 0.5 µm, with a collection efficiency, expressed by a D_50_ value, exceeding 0.5 µm. To maintain sample integrity, samples were stored in 15 ml of a nucleic acid preservation solution (reference). To minimize evaporation, we supplemented the samples with a top-off flow rate of 0.5-1.5 ml/min of molecular-grade water. Before each sampling event, we thoroughly decontaminated the Coriolis sampler as previously described ^7^.

### DNA Extraction, DNA Library Preparation, and Metagenomic Sequencing

We extracted DNA from the filters using the PowerWater DNA Isolation Kit (Qiagen, Dresden, Germany) following the manufacturer’s protocol. Shotgun metagenome libraries were generated using the Nextera XT Library Preparation Kit according to the manufacturer’s instructions (Illumina). The libraries were pooled and sequenced with paired-end 2 × 150 bp reads on an Illumina NovaSeq6000 sequencer, utilizing a high-output S4 flow cell. The library preparation and sequencing processes were conducted at the Rush Genomics and Microbiome Core Facility (GMCF) located at Rush University. Detailed information regarding this process has been previously described ^7^.

### Data Processing and Metagenomic Analysis

We performed a comprehensive quality assessment of the metagenomic datasets using FastQC software (version 0.11.9, www.bioinformatics.babraham.ac.uk/projects/fastqc). Quality trimming and Illumina adapter removal were executed using Trimmomatic (version 0.75). Applying Trimmomatic’s quality control steps resulted in an average of 4.3 million high-quality reads per sample (ranging from 3 to 11 million paired-end reads) that met the predefined quality criteria ^7^.

We initially classified the metagenome libraries into three distinct groups (Supporting Figure 1), each representing dominant atmospheric conditions and therefore the similar microbial composition ^7^. Based on this specific criterion, we then merged these groups for co-assembly. In this context, t refers to the sample identifier used in our previous study, ranging from the beginning of the sampling campaign (t1) to the end (t64): c_Northwest_1 (t12, t15, t16, t17, t23, and t24) and c_Southwest_1 (t5, t13, t14, t20, t22, t30, t31, t34, and t35) represent northwesterly and southwesterly air masses, respectively. These were observed under clear conditions before a period of consecutive dust storms and low daily mean temperatures. c_Northwest_2 (t50, t51, t56, t57, and t58) and c_Southwest_2 (t37, t38, t41, t42, t45, t49, t52, t55, t59, and t64) represent northwesterly and southwesterly air masses, respectively. These were observed under clear conditions following a period of consecutive dust storms and increasing daily mean temperatures. d_East (t1, t27, t28, t29, t32, and t62) and d_Southwest (t6, t7, t8, t9, t10, t11, t21, t33, and t36) represent dusty conditions of easterly and southwesterly air masses.

We merged the high-quality reads and subsequently assembled and binned them using the metaWRAP (v.1.3.2) pipeline ^31^. Briefly, for each sample, we co-assembled reads using the metaWRAP assembly module, which conducts the assembly with metaSPAdes (--metaspades) ^32^. We removed contigs shorter than 500 bp. To improve binning, we reassembled the reads that did not map back to the metaSPAdes assembly using MegaHit ^33^. We then binned metagenomic assemblies using three different algorithms: MaxBin2 ^34^, CONCOCT ^35^, and metaBAT2 ^36^, utilizing the Binning module (-maxbin2 -concoct -metabat2). Given the relatively low depth of these samples, we set the minimum completion to 5% and the maximum contamination to 10%. We subsequently refined each bin set using the bin_refinement module with the parameters (-c 5 -x 10). We then further improved the consolidated bin sets using the reassemble_bins module. We evaluated the completeness and contamination of bins using CheckM ^37^ within this pipeline.

We taxonomically annotated the MAGs using GTDB-Tk (v2.1.1) ^38^ with the classify workflow (classify_wf). A phylogenomic tree of these MAGs was created using the Interactive Tree of Life (v6) ^39^. Finally, we functionally annotated the MAGs using eggNOG-mapper (v2.1.12) ^40^ against the EggNOG 6.0 database ^41^ with the following parameters: --itype genome and -m diamond.

We predicted open reading frames (ORFs) for protein-coding genes using Prodigal (version 2.6.3) ^42^ with the ‘-p meta’ option, which is recommended for metagenomic datasets as it enables anonymous processing. To annotate antimicrobial resistance and virulence-related functions in ORFs, we used hmmscan (HMMER, v3.3.2-gompic-2020b) ^43^ with a stringent e-value threshold of 1×10^−5^. In instances where the same ORFs produced multiple hits to the same Pfam entry, we selected only the top matches with the highest alignment reliability (acc value), scores, and the lowest E-values. For antibiotic resistance, we utilized the Resfams HMM Database (Core, v1.2) ^44^. To identify virulence factors, we initially searched for Pfam modules related to ‘virulence’ and ‘pathogenicity’ and downloaded the corresponding Pfam profiles ^45^. We manually checked the virulence role of these Pfam modules through a literature search and removed spurious modules from our list. Subsequently, we integrated all Pfam models, including the Resfams HMM Database, and performed hmmscan as described above.

We used PLASMe ^46^ to identify plasmid contigs from metagenome assemblies, setting the minimum coverage and identity thresholds for BLASTN at 90% and 70%, respectively (-c 0.9, -i 0.7, -p 0.5). We chose PLASMe due to its effectiveness with the short contig lengths. PLASMe’s hybrid approach, which combines sequence alignment and deep learning, outperforms other methods for short contig lengths. In contrast, many widely used tools require contig lengths longer than 1000 bp for accurate prediction.

To identify putatively mobile elements from metagenome assemblies, we used several tools. We employed hmmscan to predict integrative and conjugative elements (ICEs) on ORFs using the CONJScan HMM database (v2.0.1) ^47^. In the hmmscan results, we retained only the best hits as previously described. Additionally, we used ISEScan ^48^ to predict insertion sequence (IS) elements with default parameters and IntegronFinder (version 2.0) ^49^ with the (--local-max) option to predict complete integrons or integron-associated elements, such as attC sites on assembled contigs.

We identified contigs linked to antimicrobial and virulence traits as putatively mobile if they were also linked to one of the IS elements, ICEs, and integrons. This classification is informed by the sizes of known composite transposons, which can range up to 65 Kbp (mean: 11.74 Kbp) ^50^, ICEs that range from 20 Kbp to 500 Kbp ^51,52^, and bacterial integrons, most of which are shorter than 10 Kbp ^53^. Given that most antimicrobial resistance and virulence-related contigs in our dataset did not exceed 10 Kbp (Figure 2C), we infer that if these contigs contain both antibiotic resistance or virulence genes and mobile genetic elements, they are likely associated. This approach is consistent with previous studies ^54,55^. Contigs not linked to any mobile elements or plasmids are considered chromosomal.

**Figure 2.**
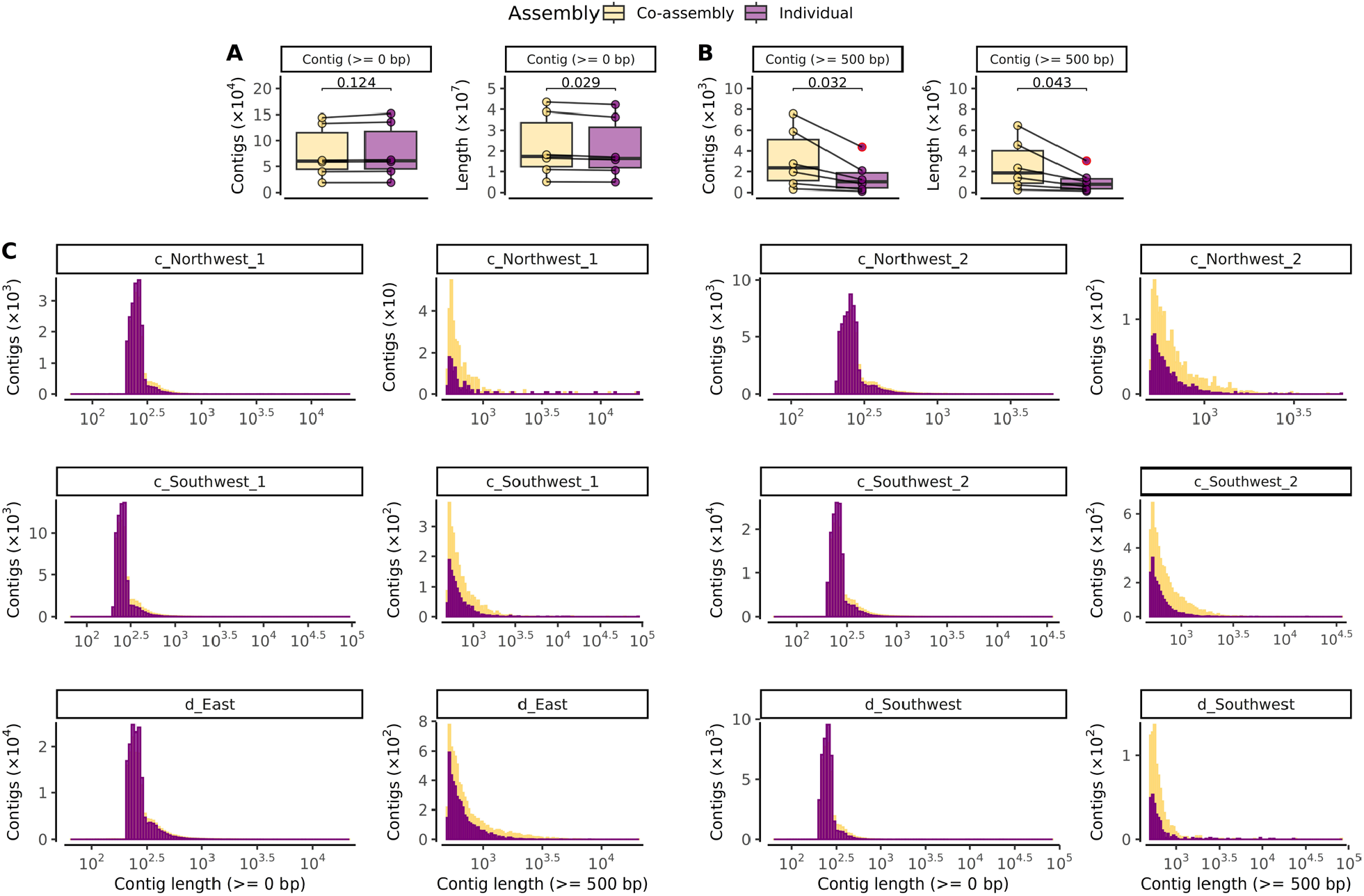
Performance comparison between co-assembly and individual assembly methods for detecting antibiotic resistance and virulence genes. (A) Total number and length of all contigs, and (B) total number and length of contigs (≥ 500 bp) annotated with at least one antibiotic resistance or virulence gene. Boxplots display median values as center lines, with lower and upper hinges representing the 25^th^ and 75^th^ percentiles. Whiskers extend from the hinges to the smallest and largest values within 1.5 times the interquartile range (IQR), and outliers are shown as red dots. Differences between the two methods were assessed using a two-sided paired t-test. (C) Length distribution of individual contigs by sample type.

To estimate the abundance, we first mapped reads from each sample to each MAG, contig, or ORF using Bowtie2 (v2.5.1) ^56^ with the (--very-sensitive-local) option. We then calculated coverage information using pileup.sh (BBMap, v39.01, sourceforge.net/projects/bbmap/) and converted it into reads per kilobase per million mapped reads (RPKM) to estimate relative abundance. This normalization procedure adjusts for both library size, accounting for the total number of mapped reads, and MAG, contig, or gene length, ensuring accurate rescaling. For ORFs predicted as partial or incomplete genes, we estimated the gene abundance accordingly.

We performed taxonomic annotation on contigs using Kraken2 (v2.1.2) ^57^ with default parameters and cross-check these results with Diamond-Megan-LR protocol ^58^, which is specifically designed for long-read sequences. Additional details on these analyses are provided in the Supporting Information.

To compare the efficiency of the co-assembly approach with assembling air samples individually, we analyzed metrics related to assembly, gene prediction, and taxonomic and functional annotation results. For this comparison, we predicted the ORFs on individual and co-assembled contigs without filtering based on assembly length. The completeness of the ORFs was defined based on Prodigal gene prediction as follows: a complete gene (Prodigal indicator “00”) has a true boundary (a start or a stop codon); a partial gene (Prodigal indicator “01” or “10”) lacks either a start or stop codon; an incomplete gene (Prodigal indicator “11”) has incomplete edges. To reduce redundancy from individual assembly results, we performed clustering on the contigs (at the nucleic acid level) and ORFs (at the amino acid level) from the individual assemblies using MMseqs2 (v13-45111) ^59^ with the following parameters: --cov-mode 1, --cluster-mode 2, -c 0.95, and --min-seq-id 0.95. Additional details on these analyses are provided in the Supporting Information. To estimate the abundance of proteins using read mapping, we used the nucleic acid sequences of the corresponding clustered proteins and calculated the RPKM as previously discussed. We then performed taxonomic and functional annotation on these non-redundant representative contigs, and genes obtained from individual assemblies and estimated the abundance as previously discussed. We evaluated the quality of individual and co-assemblies using QUAST (v5.2.0) ^60^.

To assess the presence of microbial species known for their pathogenic potential within the atmospheric microbiome, relevant to human, animal, and plant health, we utilized the extensive list provided by the Global Catalogue of Pathogens (gcPathogen) (downloaded on 4 June 2024) ^61^. This list consists of 1,317 taxa, including 510 bacterial, 407 fungal, 226 viral, and 174 parasitic taxa. We compared this list to Kraken2 and Diamond-Megan-LR results to identify the presence of these species in air samples.

### Data availability

The metagenomic sequences utilized in this study have been deposited in the National Center for Biotechnology Information (NCBI) database under the accession number PRJNA1045528.

## Results

### Metagenome Assembly

To address the challenges posed by the low-biomass nature of our samples, we grouped 45 air samples into 6 distinct subgroups based on their taxonomic and functional characteristics, as detailed in the materials and methods section. We then co-assembled these samples.

We sought to compare co-assembly with individual assembly for the same samples. As all reads are assembled together, co-assembly results in a non-redundant set of contigs and genes. In contrast, the individual assembly approach may produce overlapping contigs and genes between samples. To accurately compare the co-assembly approach to individual assembly, we reduced this redundancy by sequence clustering the contigs, and genes obtained from individual assemblies (i.e., grouping similar sequences into families), and grouping them according to the co-assembled samples. We found that the co-assembly approach yielded a higher number of longer contigs (762,369 contigs, ≥ 500 bp) and a greater total contig length (555.79 million bp in contigs ≥ 500 bp) compared to the individual assembly using the same sequencing database containing 45 air samples (455,333 and 334.31 million bp, respectively; Supporting Table 1). Specifically, co-assembly produced a significantly higher number of contigs and longer total contig length (≥ 500 bp) compared to individually assembled samples (two-sided *t*-test, *p*=0.016 and *p*=0.018 for contig number and length, Figure 1A and 1B). Not surprisingly, however, the differences in the number and total length of all contigs (≥ 0 bp) and much longer contigs (≥ 5000 bp) were not significant across all samples.

Next, we compared the co-assembly approach to individual assembly regarding their performance for gene prediction on all contigs. We found that the co-assembly approach resulted in a significantly higher number of complete genes and longer total gene lengths compared to the individual assembly approach across all samples (two-sided *t*-test, *p*=0.014 and *p*=0.015 for gene number and length, Figure 1C and 1D). However, there were no significant difference between the two methods in partial and incomplete genes, except for the number of partial genes, where the individual assembly method yielded significantly higher numbers (two-sided *t*-test, *p*=0.009). These differences between the two methods were more pronounced when their cumulative gene length distributions were analyzed (Figure 1E-G).

Furthermore, we found that the co-assembly produced a significantly higher total number (two-sided *t*-test, *p*=0.032) and length (two-sided *t*-test, *p*=0.043) of ORFs annotated with at least one antibiotic resistance or virulence gene on longer contigs (≥ 500 bp). For all contigs, these differences were only significant in total contig length (two-sided *t*-test, *p*=0.029), though the effect was less pronounced. These results are presented in Figure 2A and 2B. The differences between the two methods were more evident when analyzing their contig length distributions (Figure 2C). Moreover, we found that the composition, diversity, and richness of antibiotic resistance and virulence genes of the assembled contigs were similar for co-assembled and individually assembled communities, although the richness was slightly higher in the individual assembly approach (Supporting Figure 2).

Next, we examined whether the increase in total contig numbers and lengths affected taxonomic annotation in community analysis using Kraken2. We found that the composition, diversity, and richness of coassembled and individually assembled communities were similar. However, we observed a statistically significant increase in the richness of antibiotic resistance and virulence-associated genes in individually assembled communities, though the difference was not substantial (Supporting Figure 3).

**Figure 3.**
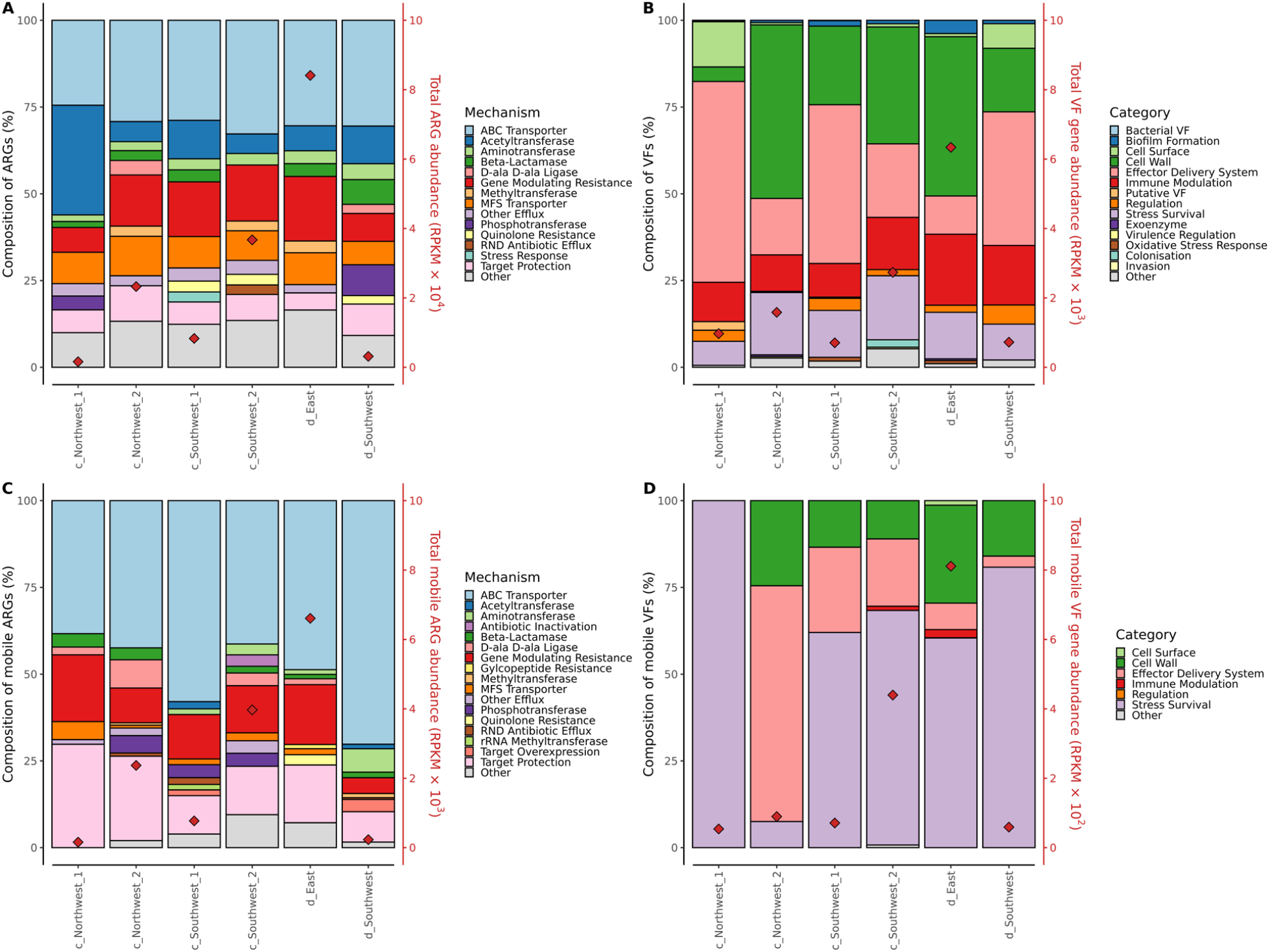
Composition of antibiotic resistance and virulence-related genes, and their predicted mobility in air samples. Composition of (A) antibiotic resistance and (B) virulence-related genes in co-assembled air samples, color-coded by antibiotic resistance mechanism and virulence category. Composition of mobile (C) antibiotic resistance and (D) virulence-related genes. Total gene abundances (RPKM) in each sample are indicated by red diamond on right y-axis.

We also verified these results using the Diamond-Megan-LR protocol to compare individual and coassembly outcomes. We found that the composition, diversity, and richness of co-assembled and individually assembled communities were similar (Supporting Figure 4). These findings aligned with the results obtained from Kraken2. However, we observed notable differences between the Kraken2 and Diamond-Megan-LR in the total number of reads mapped, and the diversity and richness of taxa across all taxonomic levels. Kraken2 showed higher diversity and richness at the species, genus, and phylum levels, although the difference in diversity at the phylum level was not significant. Moreover, the total number of reads mapped to a taxonomy varied more with each method and depended on the sample (Supporting Figure 5).

**Figure 4.**
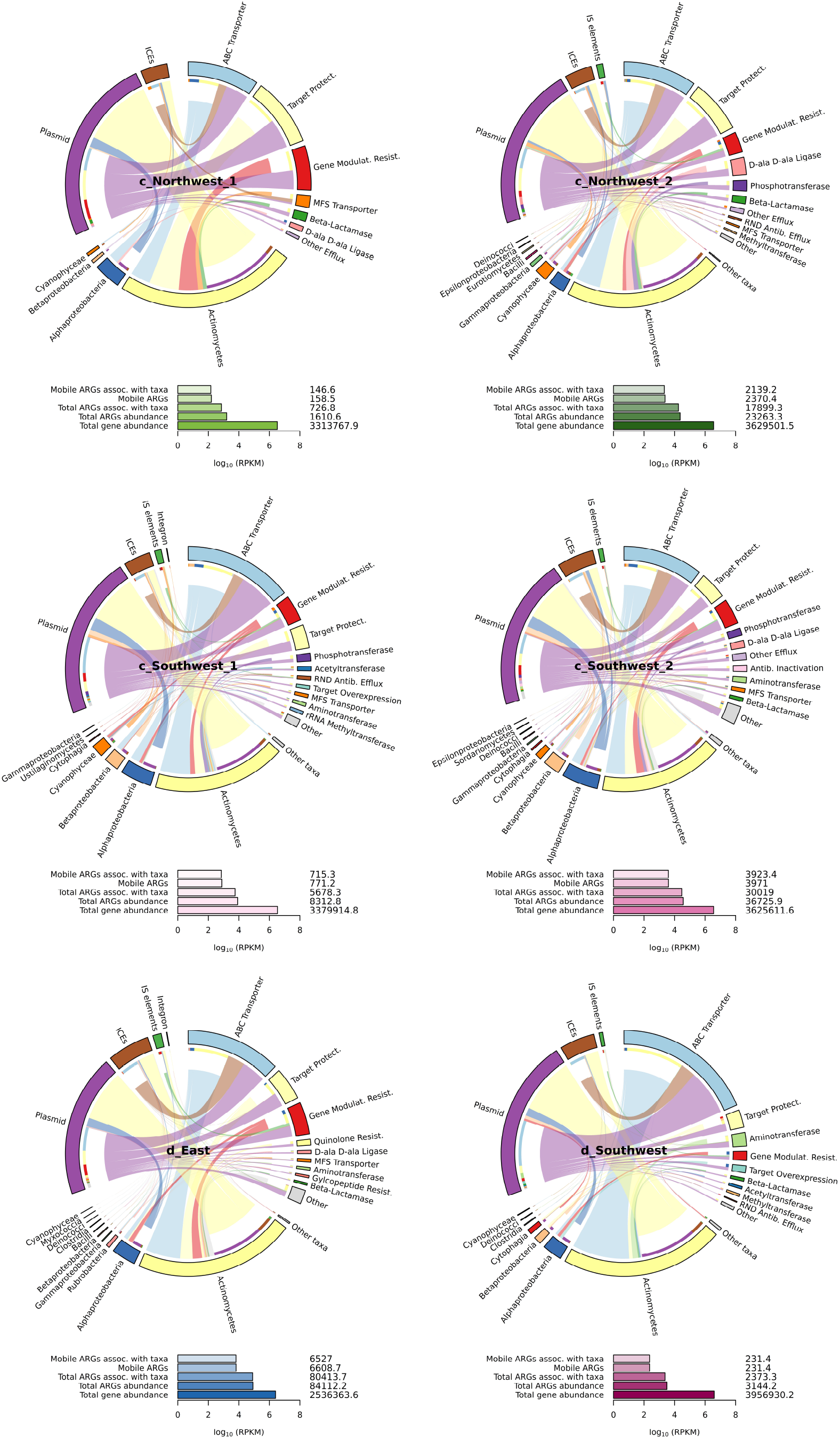
Host taxa carrying mobile antibiotic resistance genes. The figure illustrates the relationships between host taxa, antibiotic resistance genes (ARGs), and mobilization elements. Air masses are represented as follows: c_Northwest_1 and c_Southwest_1 indicate northwesterly and southwesterly air masses, respectively, observed under clear conditions before consecutive dust storms and during periods of low daily mean temperatures. c_Northwest_2 and c_Southwest_2 represent the same air masses under clear conditions following consecutive dust storms and during periods of rising daily mean temperatures. d_East and d_Southwest correspond to dusty conditions from easterly and southwesterly air masses. The “Taxa” section at the bottom of the chart displays the top 10 taxa at the class level. If more than 10 taxa are identified at the class level, or if taxa are identified at a lower taxonomic level, they are grouped as “other taxa”. The “ARG” section at the top right shows the top 10 most abundant ARGs according to mechanism they confer resistance. The “Mobility Elements” section at the top left displays the mobility elements associated with these taxa and ARGs. Connections between features are visualized, such as the association of Actinomycetes with plasmids, represented by a yellow track linking Actinomycetes to plasmids (purple), with a gap denoting the annotation transition. Bar plots at the bottom of each chord diagram present total gene (ORF) abundance, total antibiotic, and mobile antibiotic resistance gene abundances, expressed as log_10_ RPKM. The exact RPKM values are displayed above each bar.

**Figure 5.**
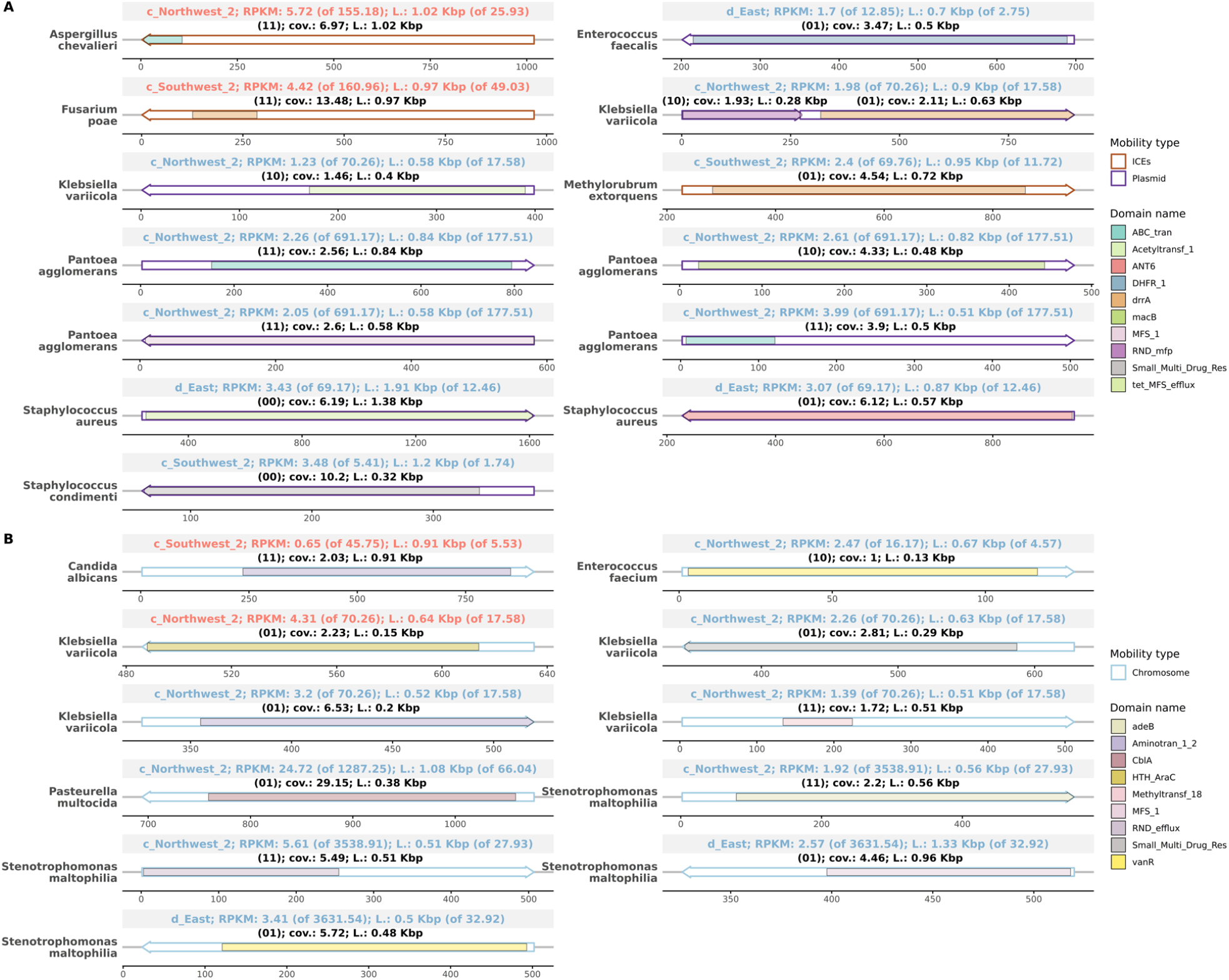
Pathogenic species carrying (A) mobile and (B) chromosomal antibiotic resistance genes. (A) DNA fragments (contigs) from pathogenic species carrying mobile antibiotic resistance genes, with only 13 contigs identified. (B) Resistance genes found on some chromosomal contigs of the most abundant pathogens linked to respiratory diseases. In both panels, the x-axis represents gene coordinates, the y-axis lists the species names associated with each contig. The direction of the arrow indicates the ORF orientation. The mobility type of each contig group is color-coded. Within each ORF, sub-gene segments indicate protein domains based on hmmscan results. These domains are aligned by gene coordinates (start and end of the domain envelope on the sequence), and each domain group is color-coded. The top gray box of each gene arrow indicates the sample name, contig abundance (RPKM), contig length, and the total contig abundance and length for contigs associated with the same species in parentheses. The text inside each box is colored according to kingdom-level taxonomy, with blue representing bacteria and red representing fungi. The completeness of the genes, indicated by the Prodigal score (01, 10, 00, and 11; see Materials and Methods) in parentheses, along with gene coverage and length on these contigs, is also provided at the top of each gene.

### Mobile antibiotic resistance genes and virulence factors

We found that a fraction of the total ORFs were annotated with antibiotic resistance and virulence-related genes across all samples (0.89±1.24% and 0.08±0.09%, mean±SD, Supporting Figure 6). Our analysis revealed that air samples contain a variety of genes associated with antibiotic resistance and virulence, with their composition and abundance varying across different conditions. Specifically, samples collected during elevated temperatures and easterly dust storms showed an increased total abundance of these genes (RPKM) compared to those collected at relatively lower temperatures, including dust storms from southwestern origins (Figure 3A and 3B). We discussed in detail the possible explanations for the differences in the total abundance of antibiotic resistance genes between atmospheric samples from different origins and conditions in a previous study ^7^.

We found that most antibiotic resistance and virulence genes were not associated with mobile genetic elements or plasmids and were therefore classified as chromosomal (90.80±1.36% and 90.70±3.76%, respectively, mean±SD). The remaining genes were associated with plasmids, ICEs, IS elements, and integrons as follows: plasmids (7.13±1.10% and 7.72±3.75%), conjugation elements (1.84±0.33% and 1.78±0.97%), insertion sequences (0.29±0.10% and 0.28±0.11%), and integrons (0.03±0.01% for antibiotic resistance contigs only). The composition of mobile antibiotic resistance and virulence genes varies widely across samples (Figure 3C and 3D). ABC Transporter genes, genes modulating resistance, and target overexpression-based antibiotic resistance mechanisms were predominant. Mobile virulence factors were less diverse, mainly dominated by functions associated with stress survival.

Acetyltransferases, which catalyze N-acetyltransferase reactions ^62^, are often found in clinical isolates where they modify antibiotics through acetylation, rendering them ineffective ^63^. Despite their overall high abundance, genes conferring antibiotic resistance through acetyltransferases were rare among mobile antibiotic resistance genes. This may be explained by the prevalence of other resistance mechanisms, such as efflux pumps, which might be favored by environmental selective pressures, resulting in less frequent mobilization of acetyltransferase genes.

The mobile genes found in air samples confer resistance against multiple antibiotics, including aminoglycosides, beta-lactams, bleomycin, fosfomycin, chloramphenicol, daunorubicin, fluoroquinolones, glycopeptides, quinolones, streptomycin, and tetracyclines, associated with mobile genetic elements (Supporting Figure 7).

### Host taxa carrying mobile antibiotic-resistance genes

We found that the most abundant taxa carrying antibiotic resistance genes are frequently associated with mobile genetic elements. However, no specific group of taxa or antibiotic resistance genes dominantly associated with mobile elements. This suggests that the observed association is primarily due to the dominance of the most abundant taxa and genes rather than any particular affinity for mobile elements. Most mobile antibiotic-resistance genes are hosted by the Terrabacteria group, particularly the Actinomycetes classes. These results are presented in Figure 4 and Supporting Table 2.

### Potential pathogens carrying antibiotic resistance genes

We found that a fraction of the total community consists of microbes known to be pathogenic to humans, animals, and plants (11.30±5.58% for Bacteria, 6.88±5.56% for Eukaryota, and 0.01±0.00% for Viruses, mean±SD, Supporting Figure 8) using Kraken2. We then cross-checked these results with those obtained from Diamond-Megan-LR protocol and observed significant differences between the two methods. While both Kraken2 and Diamond-Megan-LR identified certain pathogens, such as *Klebsiella pneumoniae, Escherichia coli, Salmonella enterica, Staphylococcus aureus*, and *Pantoea agglomerans*, their abundance estimates varied. The Megan-LR protocol identified fewer taxa overall compared to Kraken2, but it also detected taxa that Kraken2 missed, such as *Vibrio cholerae, Klebsiella oxytoca*, and *Alternaria* species (Supporting Table 3). Information on potential hosts and diseases triggered by these species is provided in Supporting Table 4.

We observed that some contigs associated with these pathogens carry antibiotic resistance and virulence genes (Supporting Table 2). Many of these contigs were chromosomal while some of them were associated with mobility elements (Figure 5). These genes are associated with factors that enhance pathogenicity in certain species but are often present in non-pathogenic microbes as well. This makes it challenging to determine whether these elements pose a potential risk. In contrast, antibiotic resistance traits in pathogenic species are likely to have more direct and significant implications in this context.

### Metagenome-assembled genomes (MAGs)

We aimed to obtain metagenome-assembled genomes (MAGs) from air samples to identify potential pathogens and to gain insight into their associations with antibiotic resistance genes. Our analysis, based on co-assembled reads, yielded only 39 MAGs. These MAGs ranged from low-quality (≤ 50% completion and ≤ 10% contamination) to medium quality (≥ 50% completion and ≤ 10% contamination) ^64^ (Supporting Table 5). Despite testing various co-assembly strategies, including combining all dusty-day samples to improve MAG completeness, we did not achieve better results. Among all the bins we created, only 13 could be phylogenetically classified (Supporting Figures 9 to 14). Clear-day air samples, influenced by northwesterly and southwesterly air masses, primarily contained MAGs of Cyanobacteriota, specifically the *Aliterella* genus, which is commonly found in aquatic environments ^65^. In contrast, dusty-day samples, influenced by easterly air masses, predominantly harbored MAGs of soil bacteria such as Actinomycetota and Bacillota, with genera including *Geodermatophilus, Kocuria, Rubrobacter* and *Arthrobacter* ^66^. These findings further validate the origin of air masses and confirms the taxonomic characterization results from our previous study using read-based methods ^7^, and the long contig-based taxonomic identifications obtained through other methods in this study. However, none of the identified MAGs belonged to known pathogenic species. This may be due to the relatively low abundance of pathogenic species, which prevented the recovery of bins that met the predesignated standards for completeness and contamination. The functional characteristics of these MAGs varied, reflecting typical microbial functions (Supporting Figure 15). However, since the MAGs were only partially assembled, estimating their full environmental potential remains a challenge.

## Discussion

In this study, we co-assembled air samples with similar air mass origins, as indicated by their comparable microbiome compositions from our previous study ^7^. We chose this method to address the challenges associated with the low biomass of atmospheric samples. While in our previous study with individual air sample assemblies, we explored the links between environmental factors and various aspects of the atmospheric microbiome robustly, it had limitations. Specifically, identifying certain features, such as genomic mobility elements, and relating these traits to functional aspects like antibiotic resistance or virulence was challenging due to the short contig lengths. Our current results suggest that co-assembling air metagenome samples of similar origins is an effective strategy for recovering more complete genes and producing longer contigs, without compromising the overall functional and taxonomic annotations of the air samples. Future studies looking to build a biome-specific gene catalog for air samples with larger datasets should consider adopting this approach ^67^. Co-assembly enabled us to generate longer contigs related to antibiotic resistance and virulence (≥ 500 bp), allowing a more accurate assessment of these features and their associations with mobility elements. However, most contigs still did not exceed 1000 bp, which is shorter compared to those found in higher biomass environments such as soil and water. To address this issue, future studies might incorporate single long DNA sequencing technologies, which could provide a promising solution. During the study, we also tested various grouping strategies for larger sample sizes, such as aggregating all samples from clear days. However, we could not assemble reads from this larger dataset, likely due to its complexity, even using a server with 750 GB of memory. This complexity likely arises from the diverse sources of emissions in the samples with limited similarity in their composition, compared to samples from a single dominating habitat, such as dust storms of specific origins.

We found a small fraction of airborne microbes as facultative pathogens carrying antibiotic resistance genes. This conclusion is based on the presence of DNA fragments from these species exhibiting these functional traits. However, interpreting these results is complex due to uncertainties about whether the DNA fragments originate from single or multiple genomes. While the co-assembly approach generally improved assembly for air samples, it still did not yield sufficiently long and informative contigs. If the DNA fragments belong to a single pathogenic species with numerous resistance genes, this could indicate a low quantity of these species but a significant risk of resistance. Conversely, suppose the fragments originate from multiple cells. In that case, it suggests a higher quantity but not multiple resistances, complicating the assessment of public health and environmental threats in this regard.

We identified a broad range of resistance mechanisms against multiple antibiotics linked to mobile genetic elements. Some airborne microbes, including certain pathogens, also carried mobile antibiotic-resistance genes, suggesting ongoing horizontal gene transfer in environmental settings. These findings are consistent with the patterns observed in the Global Microbial Gene Catalog (v1.0), which highlights the diversity of antibiotic resistance genes and their frequent association with mobile genetic elements. Notably, resistance genes and mobile elements were significantly more prevalent across diverse habitats compared to other genes ^68^. The genetic reservoir in soil suggests a continuous emergence of new antimicrobial resistance traits. These mobile resistance genes could transfer to pathogens in environmental settings or within the microbiota of humans and animals. The dispersal of these elements through air masses and dust storms further illustrates how resistance genes can spread across ecosystems, promoting their global distribution. This underscores the interconnected and global nature of antibiotic resistance. Monitoring and understanding this spread are crucial for developing strategies to mitigate its impact on public health, the environment, and agriculture.

Metagenomic analysis of air samples offers a comprehensive view of microbial dynamics and their potential implications. However, this approach has limitations and should be interpreted with caution, especially when not complemented by other methods ^69^. For instance, we identified some antibiotic resistance-related protein domains, such as PF12847 (a methyltransferase domain) ^70^ and PF02467 (a transcription factor, WhiB) ^71^, involved in essential housekeeping functions. This overlap complicates distinguishing between routine cellular processes and modifications contributing to antibiotic resistance.

Some protein domains related to antibiotic resistance, found on fungal contigs (e.g., PF00903, typically associated with glyoxalase enzymes, and fosfomycin resistance proteins like FosB ^72^, FosA ^73^, and FosX ^74^). This raises the possibility that fungi might possess genetic potential for antibiotic resistance, or that horizontal gene transfer has occurred. Given the limited understanding of antifungal resistance and the focus of resistance databases on bacteria, it remains unclear whether these findings are significant for fungal resistance or related to other cellular processes.

We occasionally observed the co-occurrence of antibiotic resistance genes and virulence factors with signatures of IS elements and ICEs within the same ORF. This dual annotation may reflect multi-domain proteins where distinct regions contribute to both resistance and mobility functions ^52^ or suggest chimeric genes formed through genetic fusion events. Annotation artifacts, particularly in complex and low-biomass datasets, may also complicate accurate identification. These scenarios highlight the need for careful consideration of genetic context and detailed analysis in metagenomic studies ^75^. The co-localization of ARGs and mobility elements can facilitate horizontal gene transfer, promoting the spread of antibiotic resistance. Future research, including experimental validation, is essential for accurately interpreting these complex associations and understanding their implications for microbial resistance and mobility in environmental contexts.

In our previous study, we found that Kraken2 is a reliable method for analyzing atmospheric communities. This was confirmed through read mapping to reference genomes, which showed reasonable and evenly distributed genome coverage, supporting the presence of species identified by Kraken2. However, when analyzing long contigs instead of reads, additional validation may be necessary. Our current study revealed instances where contigs matched specific taxa with minimal k-mer coverage. To address this, we employed the Diamond-Megan-LR protocol to cross-check Kraken2 results. While we found agreement for certain species, as detailed in the results section, we also observed significant differences between Kraken2 and Diamond-Megan-LR in atmospheric samples. These variations align with previous studies that reported diverse outcomes using different taxonomic profilers (ref). The differences likely arise from the distinct databases and methods used by the programs. Kraken2 relies on RefSeq complete genome sets, while Diamond uses the NCBI-nr database. Kraken2’s k-mer based approach identifies unique DNA segments (35 bp), whereas Diamond uses full sequence alignment to a protein reference. Kraken2’s sensitivity can lead to false positives, whereas Diamond-Megan-LR’s stringent approach may result in false negatives. These findings highlight the necessity of using multiple tools in microbiome studies, particularly for low-biomass and highly diverse environmental samples like air, which are often underrepresented in reference genome databases. Using complementary methods is essential for reliable analysis and result validation, especially in studies aimed at detecting potential pathogens and understanding their impact on public and ecosystem health.

## Author Contributions

B.A.E. conceptualized and designed the study, conducted bioinformatics and data analysis, interpreted, and visualized the results, and drafted the manuscript. Y.R. and D.Z. contributed to reviewing and editing the manuscript. All authors actively participated in result discussions and approved the final manuscript.

## References

1 Failor, K. C., Schmale, D. G., Vinatzer, B. A. & Monteil, C. L. Ice nucleation active bacteria in precipitation are genetically diverse and nucleate ice by employing different mechanisms. The ISME Journal 11, 2740–2753 (2017). 10.1038/ismej.2017.124

2 de Araujo, G. G., Rodrigues, F., Gonçalves, F. L. T. & Galante, D. Survival and ice nucleation activity of Pseudomonas syringae strains exposed to simulated high-altitude atmospheric conditions. Scientific Reports 9, 7768 (2019). 10.1038/s41598-019-44283-3

3 Bauer, H. et al. Airborne bacteria as cloud condensation nuclei. Journal of Geophysical Research: Atmospheres 108 (2003). 10.1029/2003JD003545

4 Amato, P. et al. Metatranscriptomic exploration of microbial functioning in clouds. Scientific Reports 9, 4383 (2019). 10.1038/s41598-019-41032-4

5 Vaïtilingom, M. et al. Potential impact of microbial activity on the oxidant capacity and organic carbon budget in clouds. Proceedings of the National Academy of Sciences 110, 559–564 (2013). 10.1073/pnas.1205743110

6 Erkorkmaz, B. A., Gat, D. & Rudich, Y. Aerial transport of bacteria by dust plumes in the Eastern Mediterranean revealed by complementary rRNA/rRNA-gene sequencing. Communications Earth & Environment 4, 24 (2023). 10.1038/s43247-023-00679-8

7 Erkorkmaz, B. A., Zeevi, D. & Rudich, Y. Dust storm-driven dispersal of potential pathogens and antimicrobial resistance genes in the Eastern Mediterranean. bioRxiv, 2024.2006.2024.600361 (2024). 10.1101/2024.06.24.600361

8 Drautz-Moses, D. I. et al. Vertical stratification of the air microbiome in the lower troposphere. Proceedings of the National Academy of Sciences 119, e2117293119 (2022). 10.1073/pnas.2117293119

9 Mazar, Y., Cytryn, E., Erel, Y. & Rudich, Y. Effect of Dust Storms on the Atmospheric Microbiome in the Eastern Mediterranean. Environmental Science & Technology 50, 4194–4202 (2016). 10.1021/acs.est.5b06348

10 Gat, D., Mazar, Y., Cytryn, E. & Rudich, Y. Origin-Dependent Variations in the Atmospheric Microbiome Community in Eastern Mediterranean Dust Storms. Environmental Science & Technology 51, 6709–6718 (2017). 10.1021/acs.est.7b00362

11 Mayol, E. et al. Long-range transport of airborne microbes over the global tropical and subtropical ocean. Nature Communications 8, 201 (2017). 10.1038/s41467-017-00110-9

12 Brodsky, H. et al. Assessing long-distance atmospheric transport of soilborne plant pathogens. Environ Res Lett 18 (2023). 10.1088/1748-9326/acf50c

13 Gonzalez-Martin, C., Teigell-Perez, N., Valladares, B. & Griffin, D. W. in Advances in Agronomy Vol. 127 (ed Donald Sparks) 1–41 (Academic Press, 2014).

14 Rotem, J. Sand and dust storms as factors leading to alternaria blight epidemics on potatoes and tomatoes. Agricultural Meteorology 2, 281–288 (1965). 10.1016/0002-1571(65)90014-2

15 Bai, H. et al. Airborne antibiotic resistome and microbiome in pharmaceutical factories. Environment International 186, 108639 (2024). 10.1016/j.envint.2024.108639

16 Li, X. et al. A metagenomic-based method to study hospital air dust resistome. Chemical Engineering Journal 406, 126854 (2021). 10.1016/j.cej.2020.126854

17 Wu, D. et al. Inhalable antibiotic resistomes emitted from hospitals: metagenomic insights into bacterial hosts, clinical relevance, and environmental risks. Microbiome 10, 19 (2022). 10.1186/s40168-021-01197-5

18 Zhu, G. et al. Air pollution could drive global dissemination of antibiotic resistance genes. The ISME Journal 15, 270–281 (2020). 10.1038/s41396-020-00780-2

19 Reska, T. et al. Air monitoring by nanopore sequencing. ISME Communications 4 (2024). 10.1093/ismeco/ycae099

20 Kormos, D., Lin, K., Pruden, A. & Marr, L. C. Critical review of antibiotic resistance genes in the atmosphere. Environmental Science: Processes & Impacts 24, 870–883 (2022). 10.1039/D2EM00091A

21 Luhung, I. et al. Experimental parameters defining ultra-low biomass bioaerosol analysis. npj Biofilms and Microbiomes 7, 37 (2021). 10.1038/s41522-021-00209-4

22 Luhung, I. et al. Protocol Improvements for Low Concentration DNA-Based Bioaerosol Sampling and Analysis. PLOS ONE 10, e0141158 (2015). 10.1371/journal.pone.0141158

23 Eisenhofer, R. et al. Contamination in Low Microbial Biomass Microbiome Studies: Issues and Recommendations. Trends in Microbiology 27, 105–117 (2019). 10.1016/j.tim.2018.11.003

24 Lever, M. A. et al. A modular method for the extraction of DNA and RNA, and the separation of DNA pools from diverse environmental sample types. Frontiers in Microbiology 6 (2015). 10.3389/fmicb.2015.00476

25 Bøifot, K. O., Skogan, G. & Dybwad, M. Sampling efficiency and nucleic acid stability during longterm sampling with different bioaerosol samplers. Environmental Monitoring and Assessment 196, 577 (2024). 10.1007/s10661-024-12735-7

26 Delgado, L. F. & Andersson, A. F. Evaluating metagenomic assembly approaches for biome-specific gene catalogues. Microbiome 10, 72 (2022). 10.1186/s40168-022-01259-2

27 Vosloo, S. et al. Evaluating de Novo Assembly and Binning Strategies for Time Series Drinking Water Metagenomes. Microbiol Spectr 9, e0143421 (2021). 10.1128/Spectrum.01434-21

28 Carr, V. R., Shkoporov, A., Hill, C., Mullany, P. & Moyes, D. L. Probing the Mobilome: Discoveries in the Dynamic Microbiome. Trends in Microbiology 29, 158–170 (2021). 10.1016/j.tim.2020.05.003

29 Siefert, J. L. in Horizontal Gene Transfer: Genomes in Flux (eds Maria Boekels Gogarten, Johann Peter Gogarten, & Lorraine C. Olendzenski) 13–27 (Humana Press, 2009).

30 Sun, D., Jeannot, K., Xiao, Y. & Knapp, C. W. Editorial: Horizontal Gene Transfer Mediated Bacterial Antibiotic Resistance. Frontiers in Microbiology 10 (2019). 10.3389/fmicb.2019.01933

31 Uritskiy, G. V., DiRuggiero, J. & Taylor, J. MetaWRAP—a flexible pipeline for genome-resolved metagenomic data analysis. Microbiome 6, 158 (2018). 10.1186/s40168-018-0541-1

32 Nurk, S., Meleshko, D., Korobeynikov, A. & Pevzner, P. A. metaSPAdes: a new versatile metagenomic assembler. Genome Res 27, 824–834 (2017). 10.1101/gr.213959.116

33 Li, D., Liu, C. M., Luo, R., Sadakane, K. & Lam, T. W. MEGAHIT: an ultra-fast single-node solution for large and complex metagenomics assembly via succinct de Bruijn graph. Bioinformatics 31, 1674–1676 (2015). 10.1093/bioinformatics/btv033

34 Wu, Y.-W., Simmons, B. A. & Singer, S. W. MaxBin 2.0: an automated binning algorithm to recover genomes from multiple metagenomic datasets. Bioinformatics 32, 605–607 (2015). 10.1093/bioinformatics/btv638

35 Alneberg, J. et al. Binning metagenomic contigs by coverage and composition. Nature Methods 11, 1144–1146 (2014). 10.1038/nmeth.3103

36 Kang, D. D. et al. MetaBAT 2: an adaptive binning algorithm for robust and efficient genome reconstruction from metagenome assemblies. PeerJ 7, e7359 (2019). 10.7717/peerj.7359

37 Parks, D. H., Imelfort, M., Skennerton, C. T., Hugenholtz, P. & Tyson, G. W. CheckM: assessing the quality of microbial genomes recovered from isolates, single cells, and metagenomes. Genome Res 25, 1043–1055 (2015). 10.1101/gr.186072.114

38 Chaumeil, P.-A., Mussig, A. J., Hugenholtz, P. & Parks, D. H. GTDB-Tk v2: memory friendly classification with the genome taxonomy database. Bioinformatics 38, 5315–5316 (2022). 10.1093/bioinformatics/btac672

39 Letunic, I. & Bork, P. Interactive Tree of Life (iTOL) v6: recent updates to the phylogenetic tree display and annotation tool. Nucleic Acids Research 52, W78–W82 (2024). 10.1093/nar/gkae268

40 Cantalapiedra, C. P., Hernández-Plaza, A., Letunic, I., Bork, P. & Huerta-Cepas, J. eggNOG-mapper v2: Functional Annotation, Orthology Assignments, and Domain Prediction at the Metagenomic Scale. Molecular Biology and Evolution 38, 5825–5829 (2021). 10.1093/molbev/msab293

41 Hernández-Plaza, A. et al. eggNOG 6.0: enabling comparative genomics across 12 535 organisms. Nucleic Acids Research 51, D389–D394 (2022). 10.1093/nar/gkac1022

42 Hyatt, D. et al. Prodigal: prokaryotic gene recognition and translation initiation site identification. BMC Bioinformatics 11, 119 (2010). 10.1186/1471-2105-11-119

43 Eddy, S. R. Profile hidden Markov models. Bioinformatics 14, 755–763 (1998). 10.1093/bioinformatics/14.9.755

44 Gibson, M. K., Forsberg, K. J. & Dantas, G. Improved annotation of antibiotic resistance determinants reveals microbial resistomes cluster by ecology. The ISME Journal 9, 207–216 (2015). 10.1038/ismej.2014.106

45 Finn, R. D. et al. Pfam: the protein families database. Nucleic Acids Research 42, D222–D230 (2013). 10.1093/nar/gkt1223

46 Tang, X., Shang, J., Ji, Y. & Sun, Y. PLASMe: a tool to identify PLASMid contigs from short-read assemblies using transformer. Nucleic Acids Res 51, e83 (2023). 10.1093/nar/gkad578

47 Cury, J., Abby, S. S., Doppelt-Azeroual, O., Néron, B. & Rocha, E. P. C. in Horizontal Gene Transfer: Methods and Protocols (ed Fernando de la Cruz) 265–283 (Springer US, 2020).

48 Xie, Z. & Tang, H. ISEScan: automated identification of insertion sequence elements in prokaryotic genomes. Bioinformatics 33, 3340–3347 (2017). 10.1093/bioinformatics/btx433

49 Néron, B. et al. IntegronFinder 2.0: Identification and Analysis of Integrons across Bacteria, with a Focus on Antibiotic Resistance in Klebsiella. Microorganisms 10, 700 (2022).

50 Nielsen, T. K., Browne, P. D. & Hansen, L. H. Antibiotic resistance genes are differentially mobilized according to resistance mechanism. GigaScience 11 (2022). 10.1093/gigascience/giac072

51 Johnson, C. M. & Grossman, A. D. Integrative and Conjugative Elements (ICEs): What They Do and How They Work. Annu Rev Genet 49, 577–601 (2015). 10.1146/annurev-genet-112414-055018

52 Wozniak, R. A. F. & Waldor, M. K. Integrative and conjugative elements: mosaic mobile genetic elements enabling dynamic lateral gene flow. Nature Reviews Microbiology 8, 552–563 (2010). 10.1038/nrmicro2382

53 Zhang, A. N. et al. Conserved phylogenetic distribution and limited antibiotic resistance of class 1 integrons revealed by assessing the bacterial genome and plasmid collection. Microbiome 6, 130 (2018). 10.1186/s40168-018-0516-2

54 Kent, A. G., Vill, A. C., Shi, Q., Satlin, M. J. & Brito, I. L. Widespread transfer of mobile antibiotic resistance genes within individual gut microbiomes revealed through bacterial Hi-C. Nature Communications 11, 4379 (2020). 10.1038/s41467-020-18164-7

55 Lee, K. et al. Population-level impacts of antibiotic usage on the human gut microbiome. Nature Communications 14, 1191 (2023). 10.1038/s41467-023-36633-7

56 Langmead, B. & Salzberg, S. L. Fast gapped-read alignment with Bowtie 2. Nature Methods 9, 357–359 (2012). 10.1038/nmeth.1923

57 Wood, D. E., Lu, J. & Langmead, B. Improved metagenomic analysis with Kraken 2. Genome Biology 20, 257 (2019). 10.1186/s13059-019-1891-0

58 Bagci, C., Patz, S. & Huson, D. H. DIAMOND+MEGAN: Fast and Easy Taxonomic and Functional Analysis of Short and Long Microbiome Sequences. Current Protocols 1, e59 (2021). 10.1002/cpz1.59

59 Steinegger, M. & Söding, J. MMseqs2 enables sensitive protein sequence searching for the analysis of massive data sets. Nature Biotechnology 35, 1026–1028 (2017). 10.1038/nbt.3988

60 Mikheenko, A., Prjibelski, A., Saveliev, V., Antipov, D. & Gurevich, A. Versatile genome assembly evaluation with QUAST-LG. Bioinformatics 34, i142–i150 (2018). 10.1093/bioinformatics/bty266

61 Guo, C. et al. gcPathogen: a comprehensive genomic resource of human pathogens for public health. Nucleic Acids Research 52, D714–D723 (2023). 10.1093/nar/gkad875

62 Burk, D. L., Ghuman, N., Wybenga-Groot, L. E. & Berghuis, A. M. X-ray structure of the AAC(6′)-Ii antibiotic resistance enzyme at 1.8 Å resolution; examination of oligomeric arrangements in GNAT superfamily members. Protein Science 12, 426–437 (2003). 10.1110/ps.0233503

63 Miller, G. H. et al. The most frequent aminoglycoside resistance mechanisms--changes with time and geographic area: a reflection of aminoglycoside usage patterns? Aminoglycoside Resistance Study Groups. Clin Infect Dis 24 Suppl 1, S46–62 (1997). 10.1093/clinids/24.supplement_1.s46

64 Bowers, R. M. et al. Minimum information about a single amplified genome (MISAG) and a metagenome-assembled genome (MIMAG) of bacteria and archaea. Nature Biotechnology 35, 725–731 (2017). 10.1038/nbt.3893

65 Rigonato, J. et al. Aliterella atlantica gen. nov., sp. nov., and Aliterella antarctica sp. nov., novel members of coccoid Cyanobacteria. International Journal of Systematic and Evolutionary Microbiology 66, 2853–2861 (2016). 10.1099/ijsem.0.001066

66 Battistuzzi, F. U. & Hedges, S. B. A major clade of prokaryotes with ancient adaptations to life on land. Mol Biol Evol 26, 335–343 (2009). 10.1093/molbev/msn247

67 Qin, N. et al. Longitudinal survey of microbiome associated with particulate matter in a megacity. Genome Biology 21, 55 (2020). 10.1186/s13059-020-01964-x

68 Coelho, L. P. et al. Towards the biogeography of prokaryotic genes. Nature 601, 252–256 (2022). 10.1038/s41586-021-04233-4

69 Ko, K. K. K., Chng, K. R. & Nagarajan, N. Metagenomics-enabled microbial surveillance. Nat Microbiol 7, 486–496 (2022). 10.1038/s41564-022-01089-w

70 Chow, C. S., Larnichhane, T. N. & Mahto, S. K. Expanding the nucleotide repertoire of the ribosome with post-transcriptional modifications. Acs Chem Biol 2, 610–619 (2007). 10.1021/cb7001494

71 Kormanec, J. & Homerova, D. Streptomyces aureofaciens whiB gene encoding putative transcription factor essential for differentiation. Nucleic Acids Research 21, 2512–2512 (1993). 10.1093/nar/21.10.2512

72 Cao, M., Bernat, B. A., Wang, Z., Armstrong, R. N. & Helmann, J. D. FosB, a cysteine-dependent fosfomycin resistance protein under the control of sigma(W), an extracytoplasmic-function sigma factor in Bacillus subtilis. J Bacteriol 183, 2380–2383 (2001). 10.1128/jb.183.7.2380-2383.2001

73 Beharry, Z. & Palzkill, T. Functional analysis of active site residues of the fosfomycin resistance enzyme FosA from Pseudomonas aeruginosa. J Biol Chem 280, 17786–17791 (2005). 10.1074/jbc.m501052200

74 Fillgrove, K. L., Pakhomova, S., Schaab, M. R., Newcomer, M. E. & Armstrong, R. N. Structure and mechanism of the genomically encoded fosfomycin resistance protein, FosX, from Listeria monocytogenes. Biochemistry 46, 8110–8120 (2007). 10.1021/bi700625p

75 Kerkvliet, J. J. et al. Metagenomic assembly is the main bottleneck in the identification of mobile genetic elements. PeerJ 12, e16695 (2024). 10.7717/peerj.16695

