## Supporting Information for "Metagenomic co-assembly reveals mobile antibiotic resistance genes in airborne microbiomes in the Eastern Mediterranean"

Burak A. Erkorkmaz *et al*.

**This PDF file includes:**

Supplementary Text

Supporting Figures 1 to 15

Supporting Tables 1 to 5

**Other Supporting Information for this manuscript include the following:**

Supporting Tables 1 to 5 (separate file)

**Supplementary Text**

**Materials and Methods**

**Data Processing and Metagenomic Analysis**

We performed taxonomic annotation on contigs using Kraken2 (v2.1.2) with default parameters [1]. We constructed a custom database of taxa-specific k-mers using complete genomes/proteins from diverse taxonomic groups, including archaea, bacteria, viruses, fungi, and protozoa, sourced from the NCBI RefSeq database on January 22, 2023 (kraken2-build --build). We obtained the complete lineage of taxonomy for each contig from the kingdom level to the species level using Taxonkit [2] with the taxdump file downloaded from the NCBI database on February 27, 2024.

We validated Kraken2 results using the Diamond-Megan-LR protocol [3], which is specifically designed for long-read sequences. For this analysis, we used the NCBI-nr database from June 2023. First, we aligned the assembled contigs to this protein reference database using Diamond Blastx with the following parameters: -F 15, -f 100, --range-culling, and --top 10. The --range-culling option prioritizes high-scoring alignments relative to other alignments covering the same region of the query. This prevents the common issue of multiple alignments concentrating in a small, highly conserved region. The --top 10 option reports all alignments with bit-scores within 10% of the highest score among competing alignments, ensuring a balanced representation of significant alignments across the query sequence. The -F 15 option applies a frameshift penalty of 15 to manage potential indels during translation. Next, we performed taxonomic binning of contigs using specific criteria: --longReads, --minSupport 0, --lcaCoveragePercent 51, and --topPercent 10. The --minSupport 0 option includes all potential taxonomic assignments without setting a threshold for whether a taxonomic node has been assigned enough weight. We set the --lcaCoveragePercent to 51%, meaning that at least 51% of the alignment length had to be covered for an assignment to be considered valid. We removed annotations corresponding to metazoa and viridiplantae. In both Kraken2 and Diamond-Megan-LR, we annotated contigs at the lowest taxonomic level if their total abundance was low (RPKM < 5).

To reduce redundancy from individual assembly results, we performed clustering on the contigs (at the nucleic acid level) and ORFs (at the amino acid level) from the individual assemblies using MMseqs2 (v13-45111) [4] with the following parameters: --cov-mode 1, --cluster-mode 2, -c 0.95, and --min-seq-id 0.95. This means contigs and proteins with ≥ 95% nucleic acid and amino acid identity was clustered. The --cov-mode 1 parameter allows clustering of fragmented sequences, which are common in metagenomic datasets. We ensured the longest sequence in each cluster becomes the representative sequence using using --cluster-mode 2, which helps to avoid fragmented sequences becoming representatives. This approach follows the recommendations of the MMseqs2 user guide. Clustering the contigs and ORFs occasionally resulted in duplicated names. To avoid conflicts in downstream processes, we renamed these duplicated contigs and ORFs.

### Figures

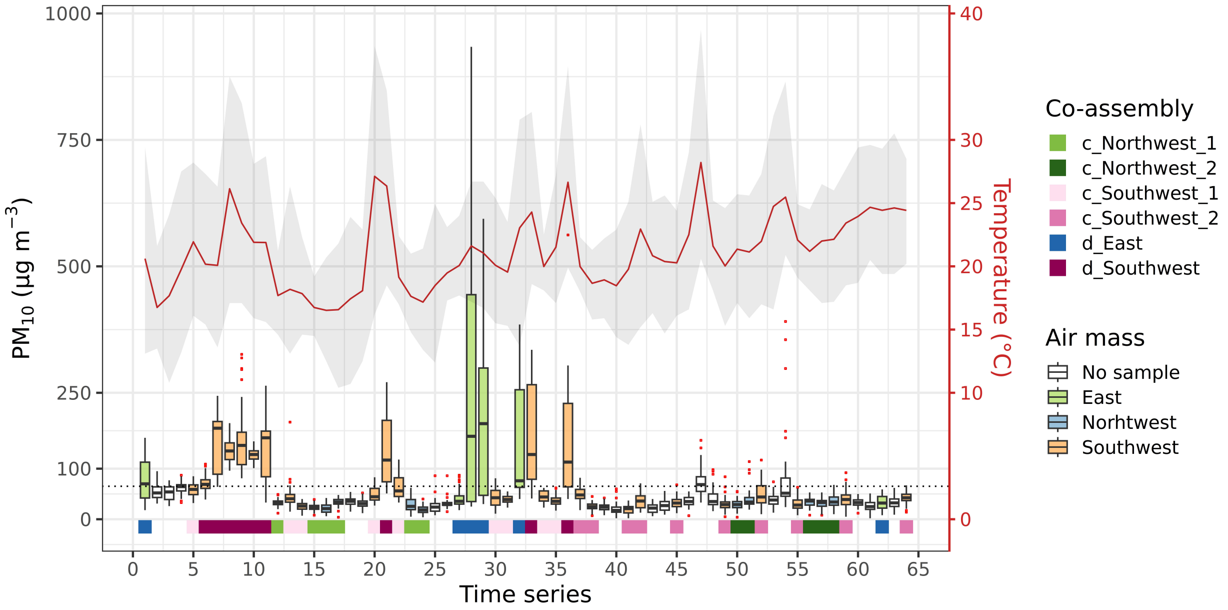


**Supporting Figure 1.** Meteorological conditions during the sampling period (March 29th to May 31st, 2022). The x-axis displays the distribution of daily $PM_{10}$ concentrations, while the y-axis represents the variations in daily temperature. The red line indicates the daily mean temperature, and the gray-colored area illustrates the range between the daily minimum and maximum temperature values. The color coding in the figure represents the air mass origins for the respective data points. Center lines in the boxplots indicate the median values, with lower and upper hinges representing the first and third quartiles (25th and 75th percentiles, respectively). Whiskers extend from the hinges to the lowest and largest values within 1.5 times the Interquartile Range (IQR) from the hinge, while outliers are depicted as red-colored dots. During the sampling campaign, white-colored box plots labeled as 'No sample' denote days when air sampling was omitted. A dotted black line within the plot signifies dust events, defined by a $PM_{10}$ concentration threshold of 65.2 µg $m^{-3}$, established based on a recent study conducted in the EM [5]. The colors below each boxplot represent co-assembled samples according to the grouping strategy described in the Materials and Methods section.

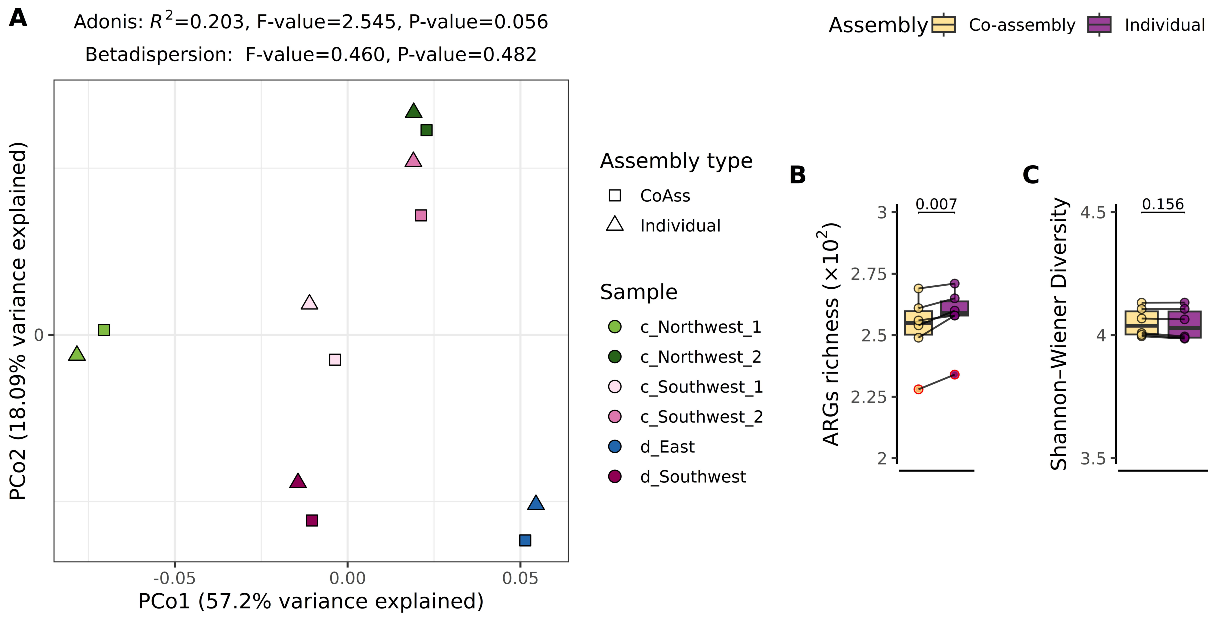


**Supporting Figure 2.** Performance comparison between co-assembly and individual assembly methods on the compositional characteristics of antibiotic resistance and virulence genes. (A) Principal Coordinate Analysis (PCoA) plot showing Bray–Curtis dissimilarity of antibiotic resistance and virulence genes based on relative abundance (calculated as RPKM values for each gene divided by the sum). The Adonis test results, which assess Bray–Curtis dissimilarity between the two methods, appear below each plot's title. Multivariate homogeneity of group dispersions (betadispersion), a prerequisite for permutational MANOVA, is also displayed. (B) Richness plot showing functional richness, representing the observed number of features. (C) Diversity plot displaying functional diversity using the Shannon–Wiener diversity index. Boxplots in (B) and (C) show median values as center lines, with lower and upper hinges representing the 25$^{th}$ and 75$^{th}$ percentiles. Whiskers extend from the hinges to the smallest and largest values within 1.5 times the interquartile range (IQR), with outliers marked as red dots. Differences between the two methods were assessed using a two-sided paired *t*-test.

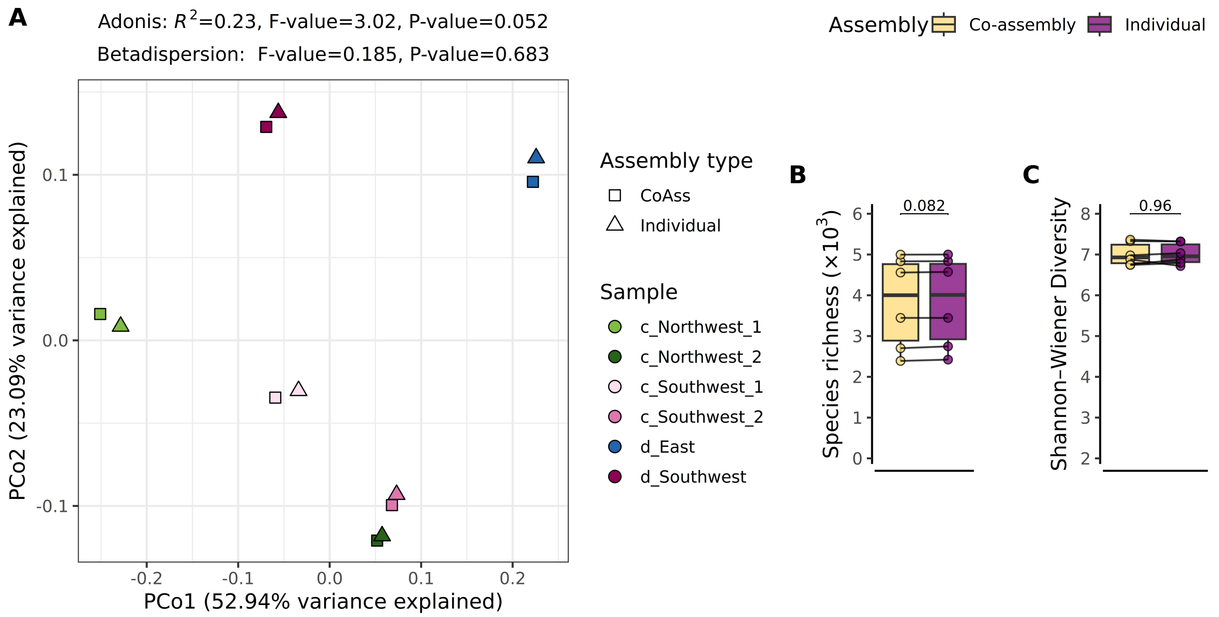


**Supporting Figure 3.** Performance comparison between co-assembly and individual assembly methods on taxonomic compositional characteristics using Kraken2. (A) Principal Coordinate Analysis (PCoA) plot showing Bray–Curtis dissimilarity of taxonomic annotation at the species level on contigs based on relative abundance (calculated as RPKM values for each contig divided by the sum). The Adonis test results, which assess Bray–Curtis dissimilarity between the two methods, appear below each plot's title. Multivariate homogeneity of group dispersions (betadispersion), a prerequisite for permutational MANOVA, is also displayed. (B) Richness plot showing taxonomic richness, representing the observed number of features. (C) Diversity plot displaying taxonomic diversity using the Shannon–Wiener diversity index. Boxplots in (B) and (C) show median values as center lines, with lower and upper hinges representing the 25$^{th}$ and 75$^{th}$ percentiles. Whiskers extend from the hinges to the smallest and largest values within 1.5 times the interquartile range (IQR), with outliers marked as red dots. Differences between the two methods were assessed using a two-sided paired *t*-test.

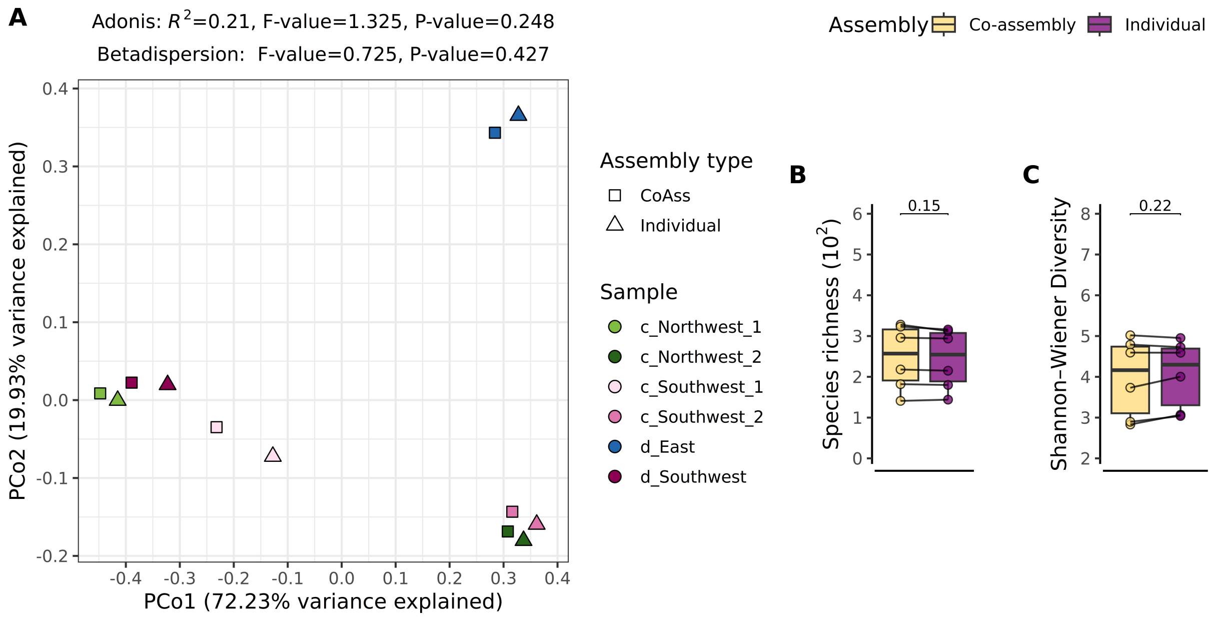


**Supporting Figure 4.** Performance comparison between co-assembly and individual assembly methods on taxonomic compositional characteristics using Diamond-Megan-LR protocol. (A) Principal Coordinate Analysis (PCoA) plot showing Bray–Curtis dissimilarity of taxonomic annotation at the species level on contigs based on relative abundance (calculated as RPKM values for each contig divided by the sum). The Adonis test results, which assess Bray–Curtis dissimilarity between the two methods, appear below each plot's title. Multivariate homogeneity of group dispersions (betadispersion), a prerequisite for permutational MANOVA, is also displayed. (B) Richness plot showing taxonomic richness, representing the observed number of features. (C) Diversity plot displaying taxonomic diversity using the Shannon–Wiener diversity index. Boxplots in (B) and (C) show median values as center lines, with lower and upper hinges representing the $25^{th}$ and $75^{th}$ percentiles. Whiskers extend from the hinges to the smallest and largest values within 1.5 times the interquartile range (IQR), with outliers marked as red dots. Differences between the two methods were assessed using a two-sided paired *t*-test.

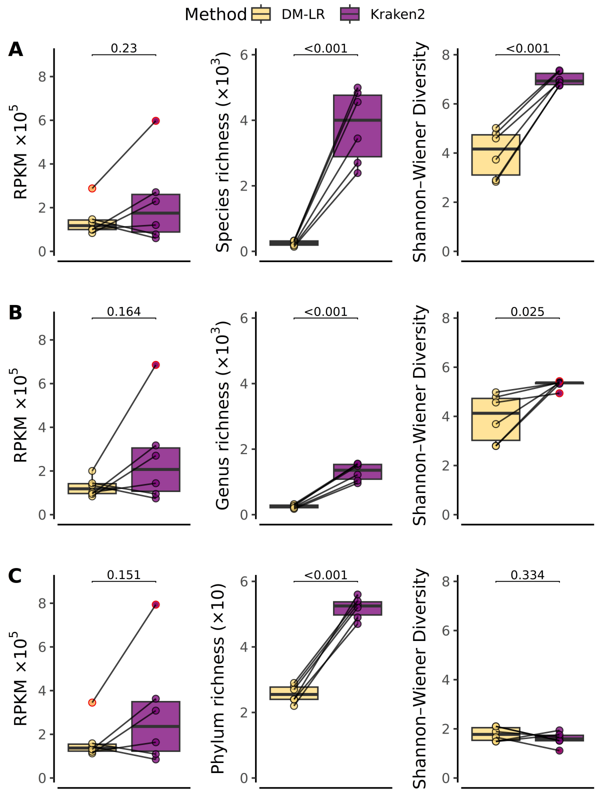


**Supporting Figure 5.** Performance comparison between the Diamond-Megan-LR protocol and Kraken2 based on total mapped reads (RPKM abundance), taxa richness, and diversity for taxonomic annotation on total contigs at (A) species, (B) genus, and (C) phylum levels. Boxplots display median values as center lines, with lower and upper hinges representing the $25^{th}$ and $75^{th}$ percentiles. Whiskers extend from the hinges to the smallest and largest values within 1.5 times the interquartile range (IQR), and outliers are marked as red dots. Differences between the two methods were assessed using a two-sided paired *t*-test.

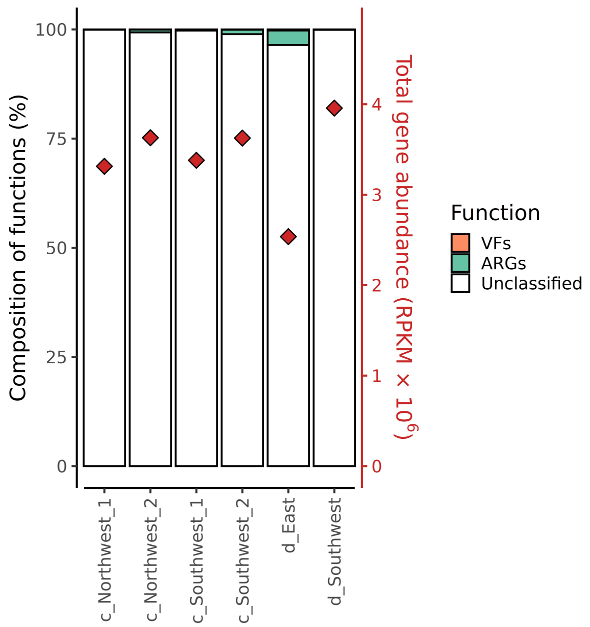


**Supporting Figure 6**. Composition of ORFs on longer contigs (≥ 500 bp) annotated with antibiotic resistance and virulence-related functions in air samples. Total ORF abundances (RPKM) for each sample are shown as red diamonds on the right y-axis. Samples are grouped and colored based on whether they exhibit antibiotic resistance or virulence traits. ORFs not annotated with these features are labeled as unclassified.

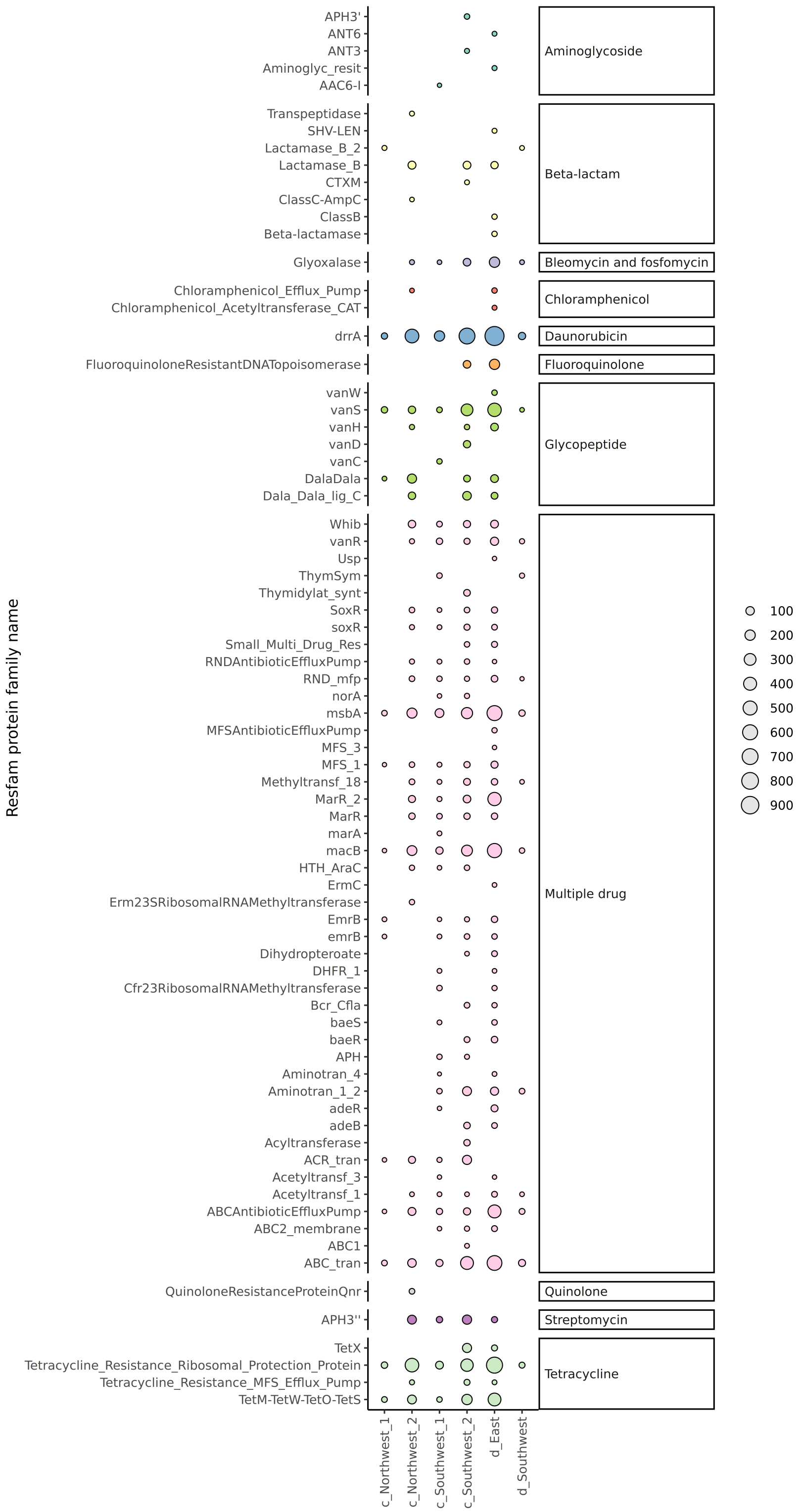


**Supporting Figure 7.** Mobile antibiotic resistance genes are annotated with protein domains based on hmmscan results from the Resfams HMM Database. Protein families are colored to indicate the specific types of antibiotics they confer resistance to and are scaled in size according to their abundance (RPKM) in each sample.

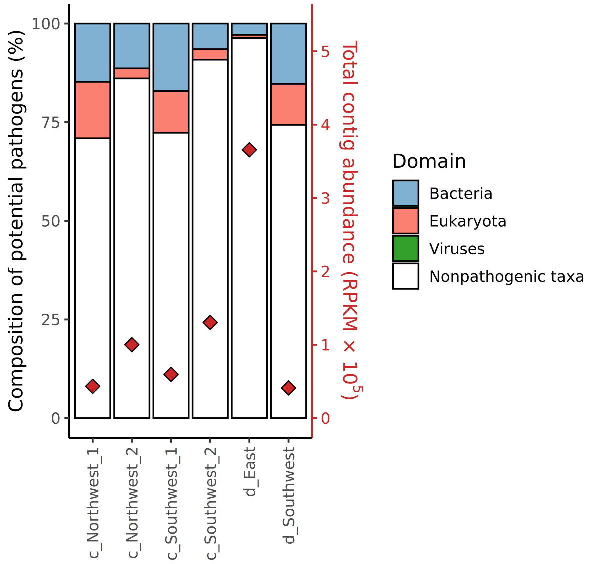


**Supporting Figure 8**. Composition of longer contigs (≥ 500 bp) annotated with taxonomy using Kraken2 in air samples. Total contig abundances (RPKM) for each sample are shown as red diamonds on the right y-axis. Samples are grouped based on their presence in the gcPathogen database and are colored according to microbial domain. Taxa not found in this list are labeled as Nonpathogenic taxa.

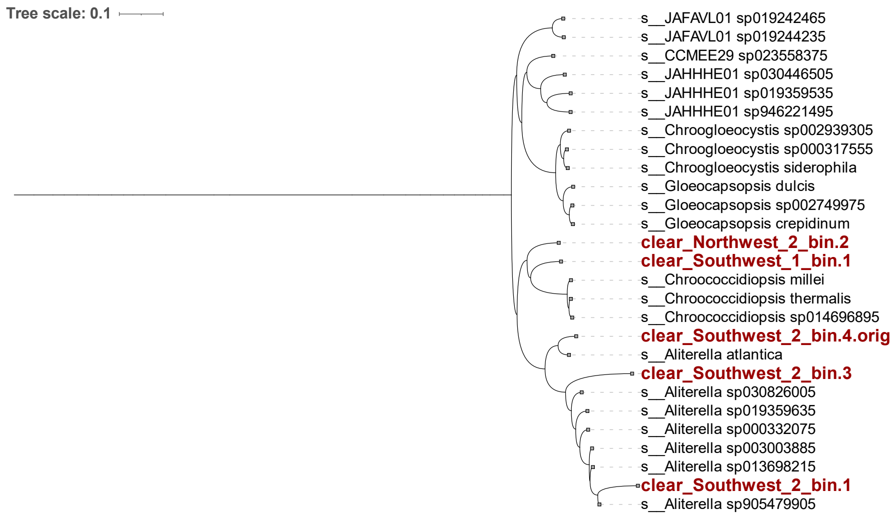


**Supporting Figure 9.** Maximum likelihood phylogenetic trees of MAGs, derived from samples collected under clear atmospheric conditions with Northwesterly and Southwesterly air masses.

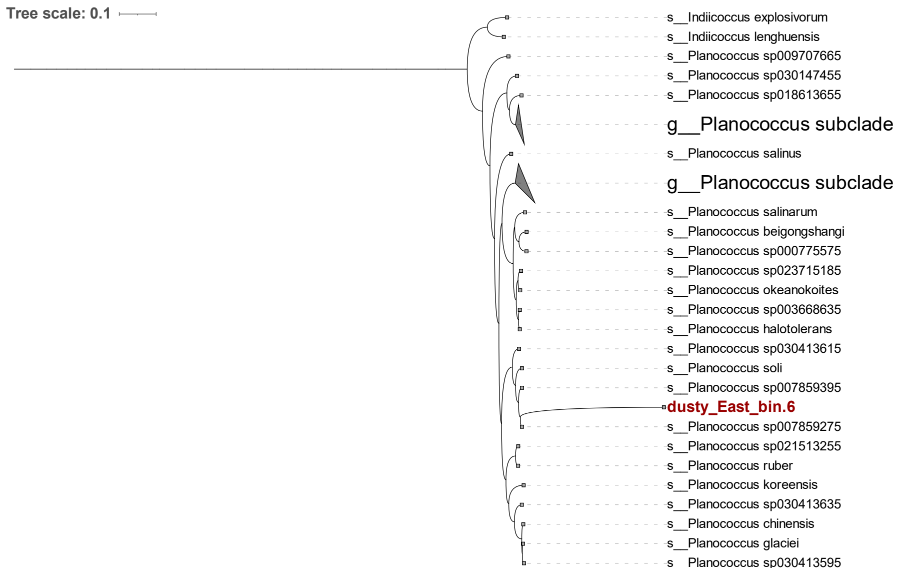


**Supporting Figure 10.** Maximum likelihood phylogenetic trees of MAGs, derived from samples collected under dusty atmospheric conditions with Easterly air masses.

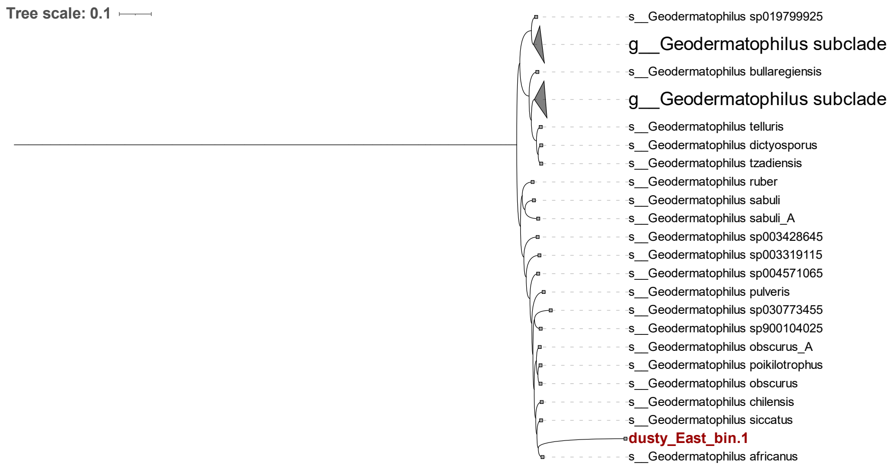


**Supporting Figure 11.** Maximum likelihood phylogenetic trees of MAGs, derived from samples collected under dusty atmospheric conditions with Easterly air masses.

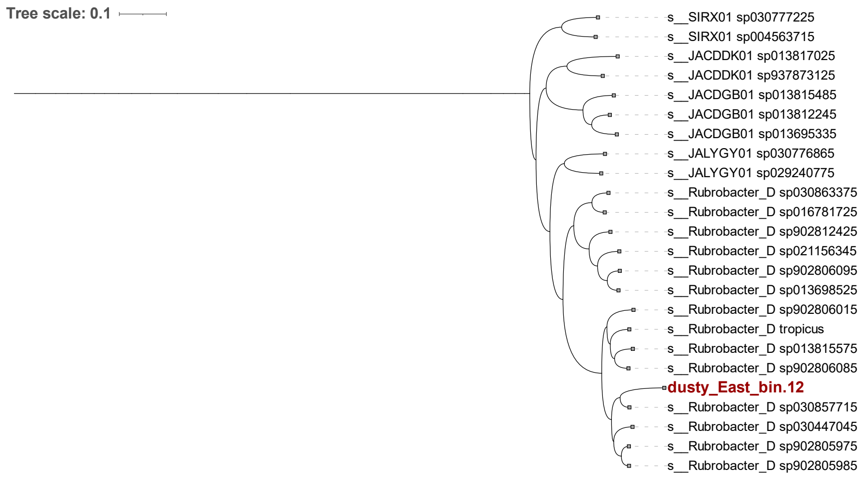


**Supporting Figure 12.** Maximum likelihood phylogenetic trees of MAGs, derived from samples collected under dusty atmospheric conditions with Easterly air masses.

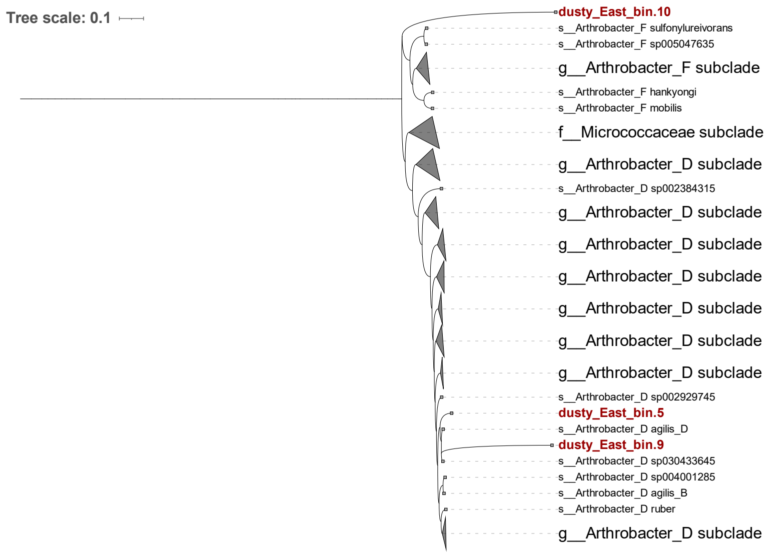


**Supporting Figure 13.** Maximum likelihood phylogenetic trees of MAGs, derived from samples collected under dusty atmospheric conditions with Easterly air masses.

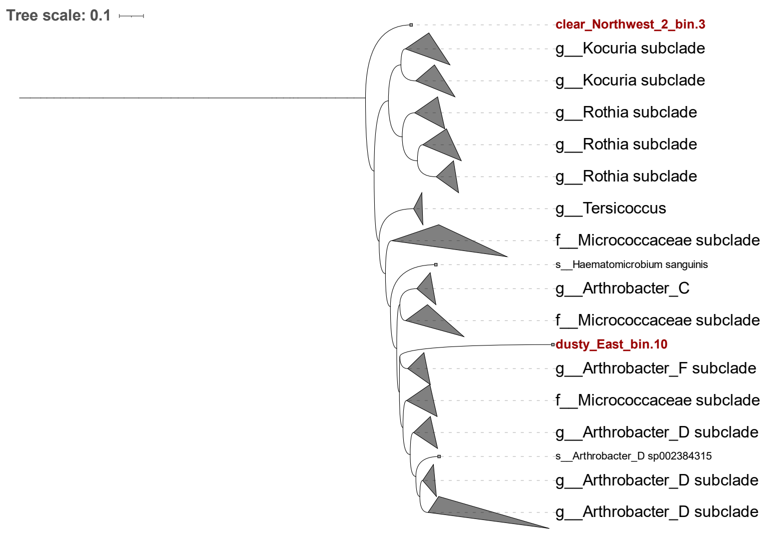


**Supporting Figure 14.** Maximum likelihood phylogenetic trees of MAGs, derived from samples collected under clear and dusty atmospheric conditions with Northwesterly and Easterly air masses.

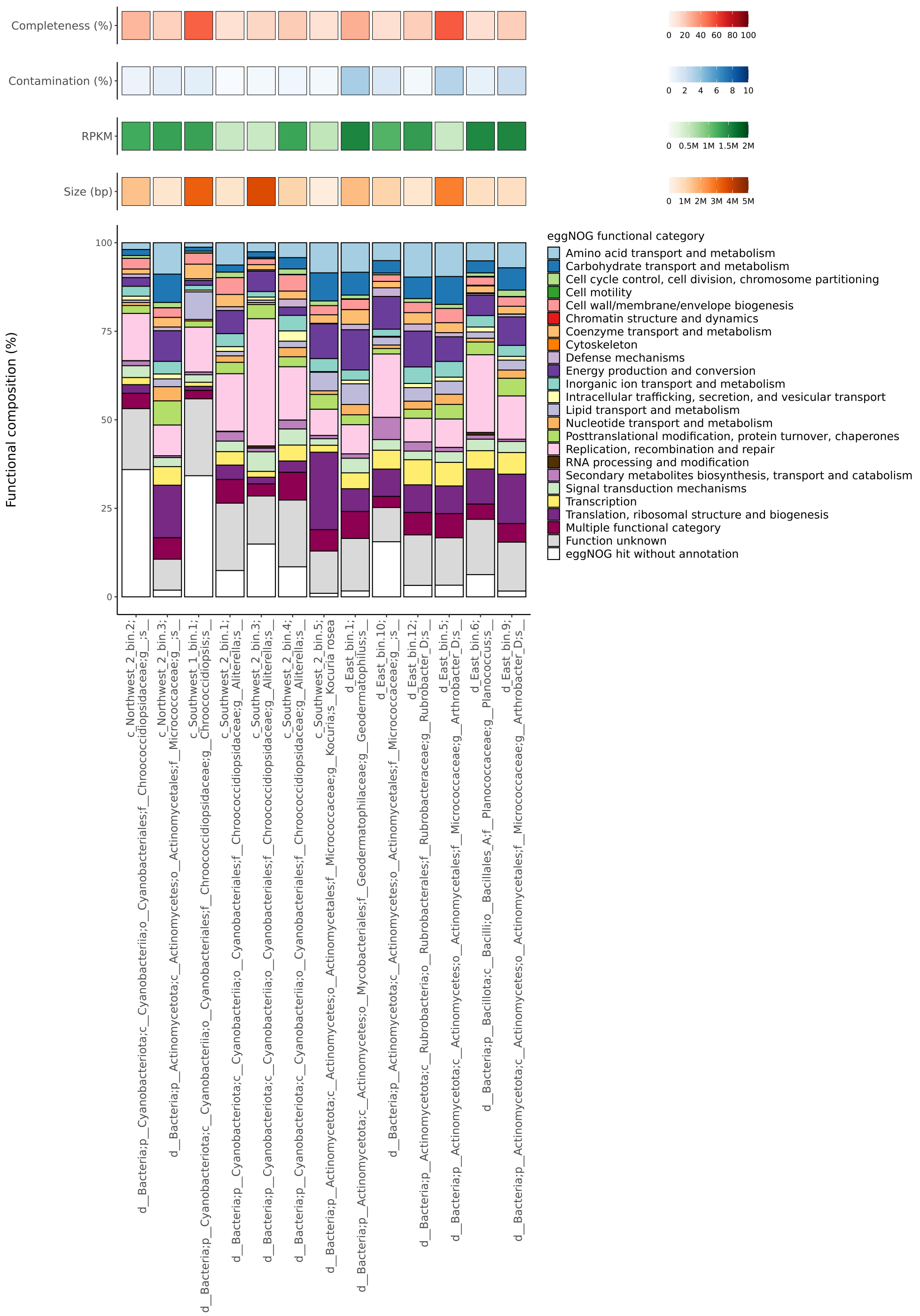


**Supporting Figure 15**. Functional characteristics of MAGs obtained from samples collected under various atmospheric conditions. Each hit from the eggNOG results is aggregated and colored according to COG functional categories. The completeness, contamination, abundance (RPKM), and size of the MAGs are displayed at the top of the bar plot. Proteins in Orthologous Groups (OG) labeled as "Function unknown" differ from "eggnog hits without annotation", which indicates either no annotation or no consensus COG for the OG. The x-axis shows the taxonomic classification based on GTDB-Tk.

**Supporting Table 1. (Separate file)** Comparison of co-assembly method performance versus individual assembly in terms of contig number and length, analysed using QUAST.

**Supporting Table 2. (Separate file)** Links microbial taxonomy to functional features on contigs, focusing on traits related to antibiotic resistance, virulence, and mobility elements such as plasmids, ICEs, IS, and integrons or integron-associated elements.

**Supporting Table 3. (Separate file)** Comparison of taxonomic abundance (RPKM) detected using Kraken2 and the Diamond-Megan-LR protocol at the phylum, genus, and species levels.

**Supporting Table 4. (Separate file)** Information on potential hosts and diseases caused by these species, based on data from the Global Catalogue of Pathogens (gcPathogen).

**Supporting Table 5. (Separate file)** CheckM results for assembled bins and taxonomic classification using GTDB-Tk.

### References

1 Wood, D. E., Lu, J. & Langmead, B. Improved metagenomic analysis with Kraken 2. *Genome Biology* **20**, 257 (2019). https://doi.org:10.1186/s13059-019-1891-0

2 Shen, W. & Ren, H. TaxonKit: A practical and efficient NCBI taxonomy toolkit. *Journal of Genetics and Genomics* **48**, 844-850 (2021). https://doi.org:https://doi.org/10.1016/j.jgg.2021.03.006

3 Bağcı, C., Patz, S. & Huson, D. H. DIAMOND+MEGAN: Fast and Easy Taxonomic and Functional Analysis of Short and Long Microbiome Sequences. *Current Protocols* **1**, e59 (2021). https://doi.org:https://doi.org/10.1002/cpz1.59

4 Steinegger, M. & Söding, J. MMseqs2 enables sensitive protein sequence searching for the analysis of massive data sets. *Nature Biotechnology* **35**, 1026-1028 (2017). https://doi.org:10.1038/nbt.3988

5 Sarafian, R., Nissenbaum, D., Raveh-Rubin, S., Agrawal, V. & Rudich, Y. Deep multi-task learning for early warnings of dust events implemented for the Middle East. *npj Climate and Atmospheric Science* **6**, 23 (2023). https://doi.org:10.1038/s41612-023-00348-9
